# Polychromatic solar energy conversion in pigment-protein chimeras that unite the two kingdoms of (bacterio)chlorophyll-based photosynthesis

**DOI:** 10.1101/565283

**Authors:** Juntai Liu, Vincent M. Friebe, Raoul N. Frese, Michael R. Jones

## Abstract

Natural photosynthesis can be divided between the chlorophyll-containing plants, algae and cyanobacteria that make up the oxygenic phototrophs and a diversity of bacteriochlorophyll-containing bacteria that make up the anoxygenic phototrophs. Photosynthetic light harvesting and reaction centre proteins from both groups of organisms have been exploited in a wide range of biohybrid devices for solar energy conversion, solar fuel synthesis and a variety of sensing technologies, but the energy harvesting abilities of these devices are limited by each protein’s individual palette of (bacterio)chlorophyll, carotenoid and bilin pigments. In this work we demonstrate a range of genetically-encoded, self-assembling photosystems in which recombinant plant light harvesting complexes are covalently locked with reaction centres from a purple photosynthetic bacterium, producing macromolecular chimeras that display mechanisms of polychromatic solar energy harvesting and conversion not present in natural systems. Our findings illustrate the power of a synthetic biology approach in which bottom-up construction of a novel photosystem using naturally disparate but mechanistically complementary components is achieved in a predictable fashion through the genetic encoding of adaptable, plug-and-play covalent interfaces.

**ToC image:** 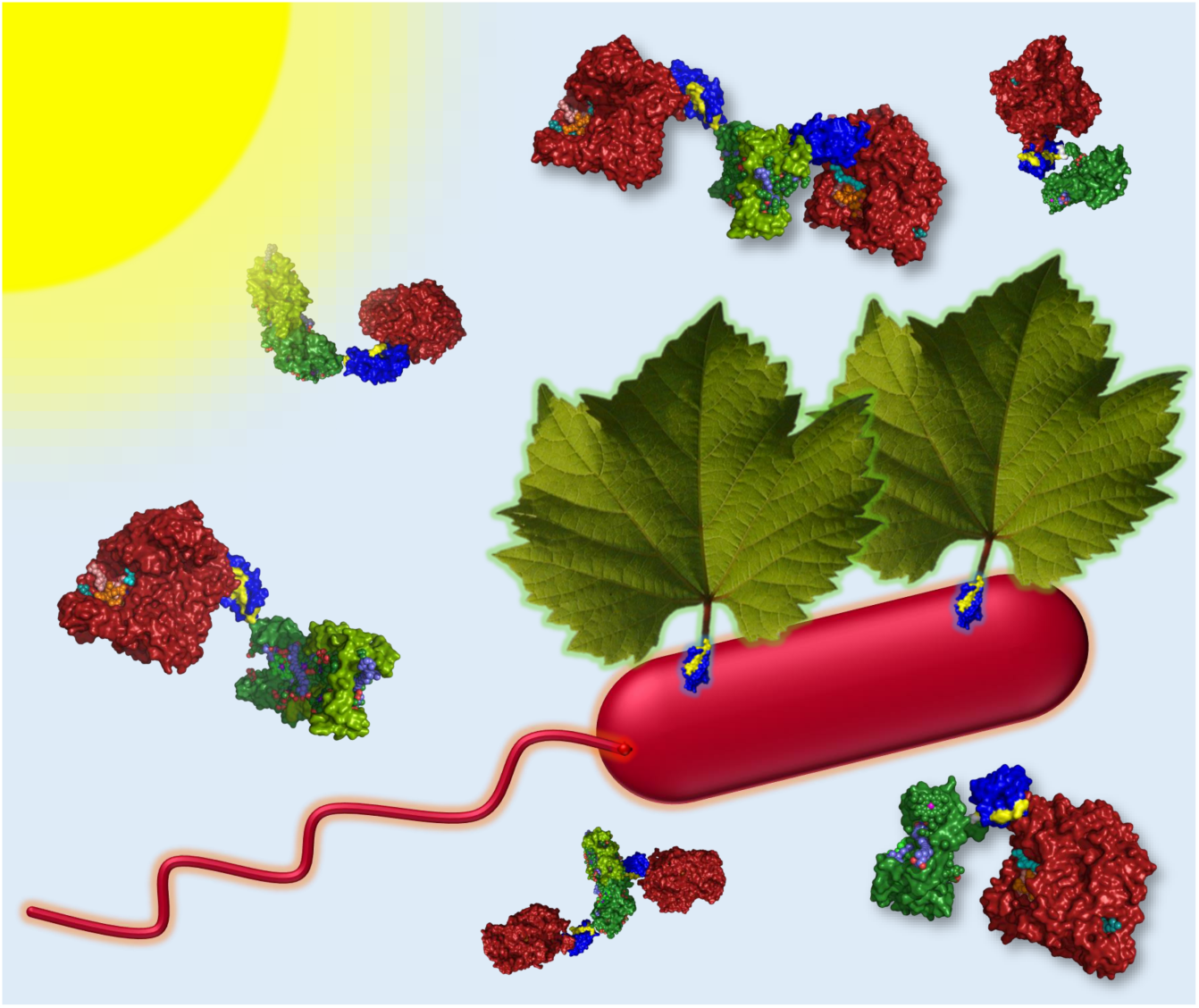

## Main

Our everyday experience of photosynthesis is dominated by the blue/red-absorbing pigment chlorophyll, a magnesium tetrapyrrole that acts as both a harvester of solar energy and a carrier of electrons and holes. Variants of this versatile molecule, principally chlorophyll *a* and chlorophyll *b*, are found in the plants, algae and cyanobacteria that make up the oxygenic phototrophs. Less well-known are the anoxygenic phototrophs, bacteria that use electron donors other than water and have one or more variants of bacteriochlorophyll as their principal photosynthetic pigment. Although these bacteria are less obvious in our environment, oxygen-tolerant species are widespread in oceanic surface waters where they make a sizeable contribution to global solar energy conversion^1^. A few species, including the bacteriochlorophyll *a*-containing *Rhodobacter* (*Rba*.) *sphaeroides*, have played major roles in our understanding of excitation energy transfer in light harvesting “antenna” complexes (LHCs)^2–4^ and charge separation in photochemical reaction centres (RCs)^5,6^.

Improving the performance of photosynthesis and finding new ways to exploit natural solar energy conversion have become important research topics^7,8^, and there is growing interest in the use of photosynthetic proteins as environmentally-benign components in biohybrid devices for solar energy conversion^9–14^. Photoexcitation of a RC in such a device triggers intra-protein charge separation, producing a potential difference between opposite “poles” of the protein that drives subsequent electron transfer to create a photocurrent and photovoltage. In addition to solar energy conversion *per se*, proposed applications of photoprotein devices have included biosensing, light/UV sensing, touch sensing and solar fuel synthesis^9–16^. Photosynthetic proteins are attractive as device components because they are environmentally sustainable and benign, they achieve solar energy conversion with a very high quantum efficiency (charges separated per photon absorbed), and they can be adapted to purpose through protein engineering. However, a limitation is their selective use of available solar energy^7,8^, a consequence of their particular palette of light harvesting pigments (Fig. 1a). This can be evidenced in devices through the recording of action spectra of external quantum efficiency (EQE – the number of charges transferred per incident photon), which exhibit peaks and troughs that correspond to the absorbance spectra of the particular light harvesting pigments that are coupled to charge separation in the device^12,17–21^.

**Fig. 1.**
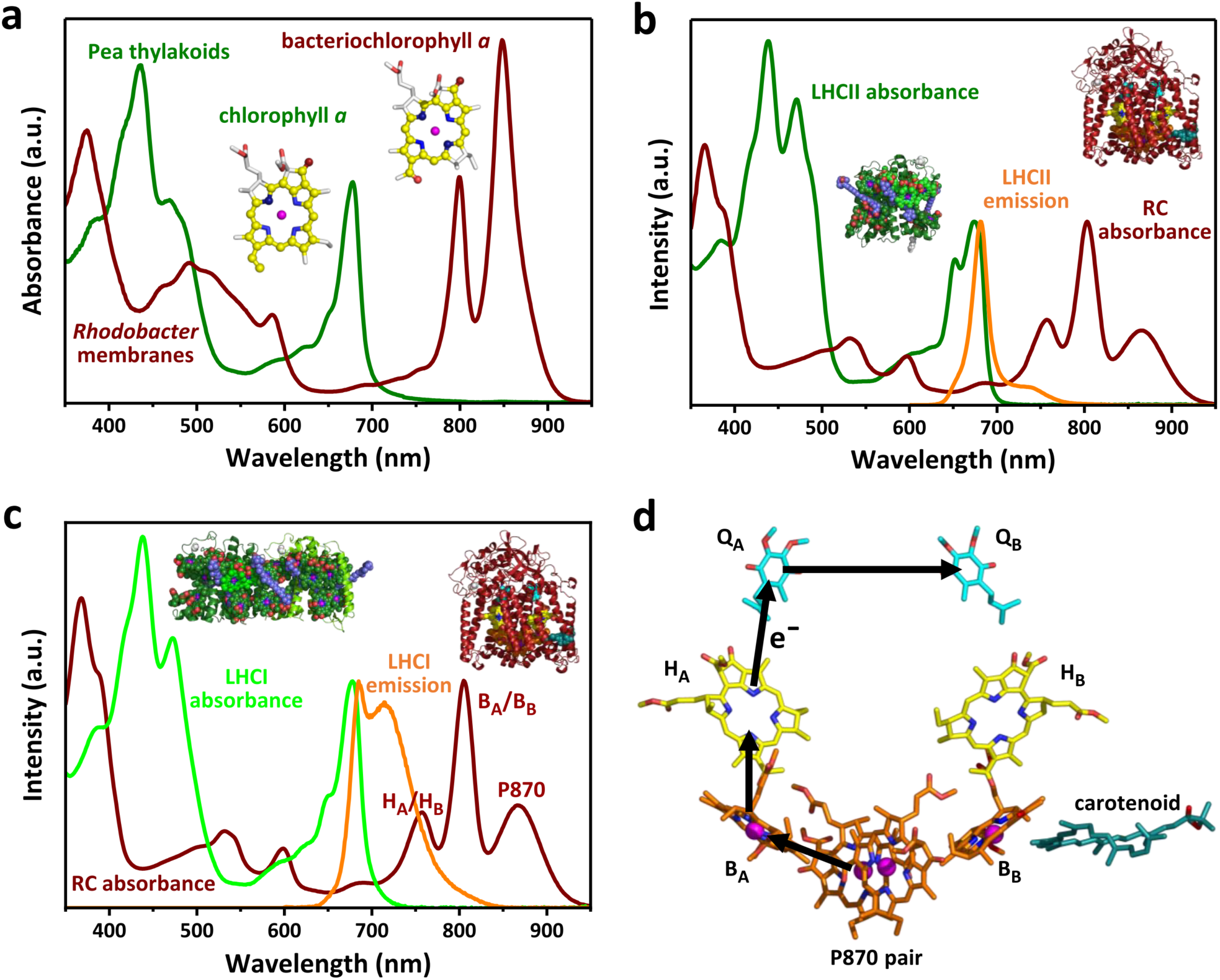
Component absorbance, emission and mechanism. **a** Thylakoid membranes from oxygenic phototrophs such as pea and chromatophore membranes from anoxygenic phototrophs such as *Rba. sphaeroides* have complementary absorbance spectra due to differences in the electronic structures of the macrocycle π electron systems of chlorophyll and bacteriochlorophyll (see also Supplementary Fig. S1). **b** The major plant light harvesting complex LHCII harvests solar energy in regions where absorbance by *Rba. sphaeroides* RCs is weak, notably around 650 nm, and its emission spectrum overlaps the absorbance spectrum of the RC between 640 nm and 800 nm. **c** The red-enhanced emission spectrum of heterodimeric plant LHCI has a stronger overlap with the absorbance spectrum of the *Rba. sphaeroides* RC, particularly the coincident absorbance bands of the bacteriopheophytins (H_A_/H_B_). **d** Architecture of the RC cofactors and the route of four-step charge separation which oxidises P870 and reduces Q_B_. The bacteriochlorophylls (orange carbons) and bacteriopheophytins (yellow carbons) give rise to the absorbance bands labelled in **c**. Further descriptions of pigment-protein structures and their sources are given in Supplementary Fig. S1.

One option for the expansion of a protein’s light harvesting capacity is to attach to it chromophores such as synthetic dyes^22–24^ or emissive nanoparticles^25–27^. Drawbacks of this approach are that synthetic dyes are often expensive and prone to photobleaching^26^, whilst fluorescent nanoparticles can be toxic and achieving well-controlled assembly of protein-nanoparticle conjugates is challenging^28^. More akin to the present study is a report of a fusion protein between a single Yellow Fluorescent Protein (YFP) and the purple bacterial RC, which has the effect of somewhat enhancing light harvesting in a region where RC absorbance is weak by adding a single chromophore^29^.

A striking observation is the complementary nature of the absorbance spectra of chlorophyll and bacteriochlorophyll photosystems (Fig. 1a). This is enabled by the somewhat different electronic structures of their principal pigments (Supplementary Fig. S1a) and facilitates the occupancy of complementary ecological niches by oxygenic and anoxygenic phototrophs. Chlorophyll absorbs most strongly in the blue and red whereas the absorbance of bacteriochlorophyll is shifted to the near-ultraviolet and near-infrared. The absorbance spectra of plant and bacterial carotenoids between 400 and 600 nm are also somewhat complementary (Fig. 1a). Thus, anoxygenic phototrophs harvest parts of the solar spectrum which oxygenic phototrophs do not absorb well, and *vice versa*.

Following nature’s lead, here we present the use of genetic encoding to achieve *in vitro* self-assembly, from diverse components (Fig. 1b,c), of novel photoprotein “chimeras” that display polychromatic solar energy harvesting and conversion. The components are the *Rba. sphaeroides* RC^5,6^ and the LHCII^30–33^ and heterodimeric LHCI^34–38^ proteins from *Arabidopsis* (*A*.) *thaliana* (Supplementary Fig. S1b-e). Highly specific and programmable self-assembly is achieved through adaptation of these components with the constituents of a two-component protein interface domain (Supplementary Fig. S1f) that covalently locks together two photosynthetic membrane proteins that have no natural propensity to associate in a specific and/or controllable manner. The resulting macromolecular, adaptable chimeric photosystems have defined compositions, and display novel mechanisms of solar energy conversion across the near-UV, visible and near-IR.

## Results

### Solar energy conversion by unadapted photosystem components

We first looked at whether plant LHCIIs can pass harvested energy to purple bacterial RCs in dilute solution in the absence of complementary genetic adaptations to promote specific heterodimerisation (complexes defined as “unadapted”). On receipt of excitation energy, photochemical charge separation in the *Rba. sphaeroides* RC is a rapid four-step process (Fig. 1d) that produces a metastable oxidised primary electron donor (P870^+^) and reduced acceptor ubiquinone (Q_B_^-^); energy transfer can therefore be detected as a quenching of LHC emission accompanied by an enhancement of P870 oxidation. Although bacterial RCs and plant LHCIIs (see Methods for sources) have overlapping absorbance and emission spectra between 640 nm and 800 nm (Fig. 1b) no appreciable energy transfer was observed when wild-type (WT) RCs were mixed in solution with an LHCII because they have no capacity for binding to one another. The addition of purified wild-type (WT) RCs did not significantly reduce emission from LHCII (Fig. 2a) and photo-oxidative bleaching of the absorbance band of this RC’s P870 primary electron donor BChls in response to 650 nm excitation was not significantly enhanced by the addition of LHCII (Fig. 2b), which absorbs strongly at this wavelength (Fig. 1b).

**Fig. 2.**
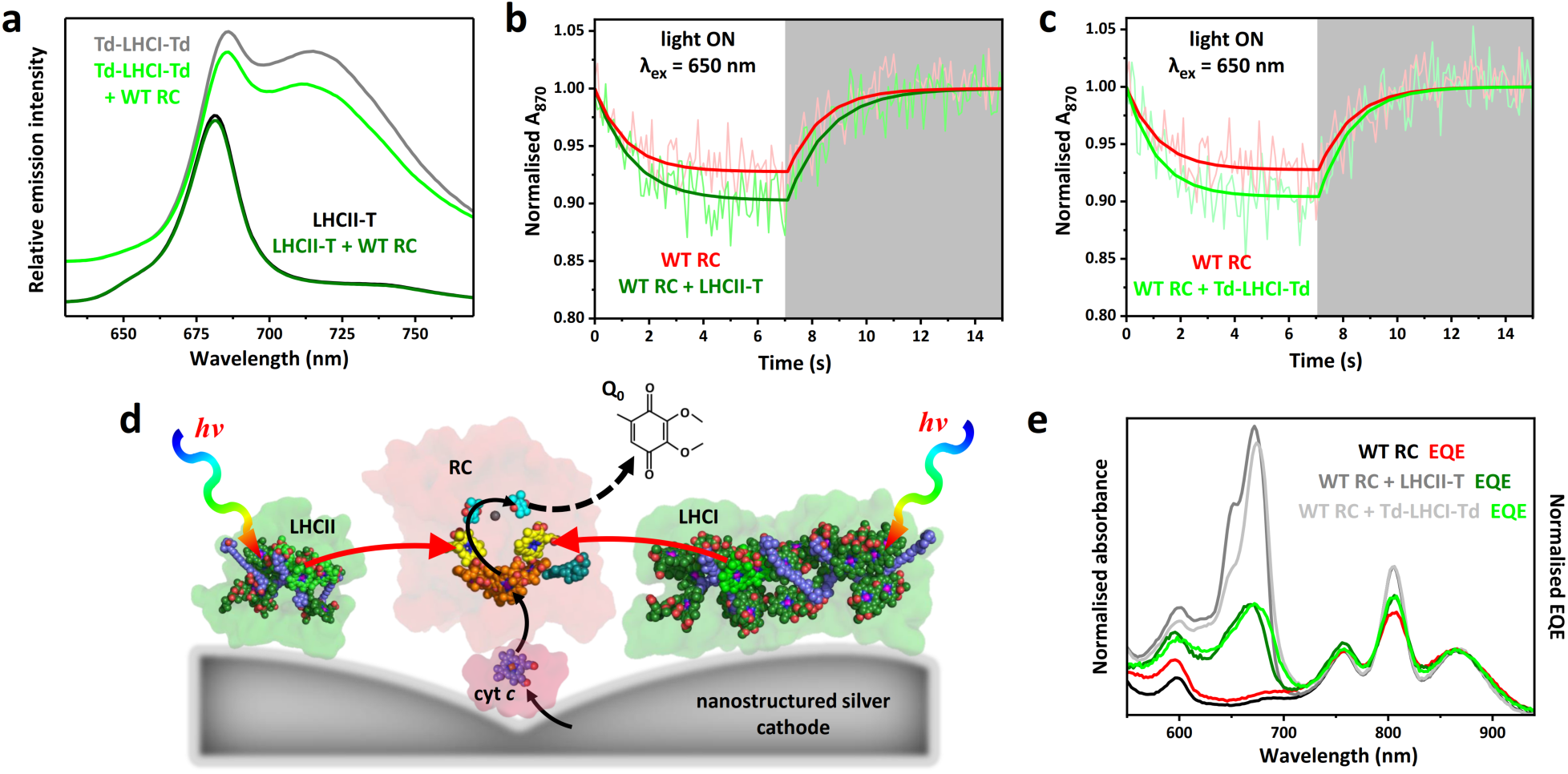
Energy transfer requires colocalization of RCs and LHCs. **a** LHCII emission (excitation at 475 nm) and LHCI emission (excitation at 500 nm) in the absence and presence of WT RCs. The latter spectra are offset for clarity. **b** Data and fits for photobleaching and dark recovery of P870 absorbance for the WT RC in the absence and presence of LHCII (using variant LHCII-T). **c** Photobleaching and dark recovery of P870 absorbance in WT RCs in the absence and presence of LHCI (using variant Td-LHCI-Td). **d** Schematic of photocurrent generation on a nanostructured silver electrode; black arrows show the route of electron transfer, red arrows show energy flow. **e** Solution absorbance and EQE spectra for WT RCs compared with those for mixtures of WT RCs with either LHCII-T or Td-LHCI-Td. The absorbance spectra were normalised at 804 nm, whilst each EQE spectrum was normalised to the corresponding absorbance spectrum at the maximum of the long-wavelength P870 band.

In comparison to LHCII, the spectral overlap (*J*) between LHC emission and RC absorbance is ∼80 % larger in the case of LHCI (Fig. 1c and Supplementary Table S1) which contains a pair of “red-form” chlorophyll *a* that possess a charge-transfer state that mixes with the low energy exciton state^38^. Although the addition of WT RCs did bring about a decrease in LHCI emission (Fig. 2a) there was no associated significant increase in RC P870 photobleaching in the presence of LHCI (Fig. 2c), leading to the conclusion that the observed emission quenching was not due to energy transfer. Protein concentrations used for the fluorescence measurements were too low (max absorbance < 0.07) for this drop in LHCI emission to be attributable to reabsorption by the added RCs, and an equivalent drop was not seen for LHCII and WT RCs at similar concentrations (Fig. 2a). As it is known that the emission quantum yield of LHCI *in vitro* is much more sensitive to its environment than is the case for LHCII^36^, the observed drop in LHCI emission on adding WT RCs is attributed to a change in its intrinsic quantum yield rather than being a signature of energy transfer.

Although no significant energy transfer was seen between these proteins in dilute solution, to establish the principle that plant LHCs can pass energy to bacterial RCs when brought sufficiently close together mixtures of LHC and WT RC proteins were adhered to a nanostructured silver cathode and their capacity for generating photocurrents examined (see Materials and Methods). In this photoelectrochemical system (Fig. 2d) cytochrome *c* (cyt *c*) is used to “wire” charge separation in the RC to the cathode, and ubiquinone-0 (Q_0_) shuttles electrons to the counter electrode^20,39,40^. Electrodes drop-cast with purified WT RCs produced a photocurrent in response to RC-specific 870 nm light and a weaker current in response to 650 nm excitation where RC absorbance is very low (Supplementary Fig. S2a). An external quantum efficiency (EQE) action spectrum showed good correspondence with the RC absorbance spectrum, confirming that the photocurrent was attributable to light capture by the pigments of the RC (Fig. 2e, black versus red). As expected, an electrode fabricated with purified LHCII failed to show any photocurrent response during 650 nm excitation of the main low-energy LHCII absorbance band (Supplementary Fig. S2a).

For electrodes fabricated from mixtures of WT RCs and LHCs, in addition to the expected RC bands the EQE spectra contained a component between 620 and 700 nm that corresponded to the low-energy absorbance band of LHCII or LHCI (Fig. 2e, green and full spectra can be found in Supplementary Fig. S3a). A contribution from the high energy Soret absorbance band of LHCII or LHCI was also observed in EQE spectra (Supplementary Fig. S4b,e,f). This demonstrated that bacteriochlorophyll-based purple bacterial RCs can utilize chlorophyll-based plant LHCs for energy harvesting, producing charge separation and a photocurrent response, provided they are brought within Förster resonance energy transfer (FRET) distance of one another. In this case this was realised by colocalising the two proteins on the surface of a biophotoelectrode.

### Design and production of component for chimeric photosystems

In an attempt to activate chlorophyll to bacteriochlorophyll energy transfer in dilute solution, RCs and LHCs were adapted using the SpyTag/SpyCatcher protein fusion system^41^ as a programmable interface. When mixed in solution, highly-specific binding of the short SpyTag peptide to the SpyCatcher protein domain initiates autocatalysis of an isopeptide bond between the two involving aspartate and lysine residues (Supplementary Fig. S1f), producing a single, covalently-locked, water-soluble protein domain^41^.

To adapt the RC for LHC binding an optimized version of SpyCatcher^42^, 106 amino acids in length (SpyCatcherΔ), was attached to the N-terminus of the RC PufL protein either directly (dubbed “RC*C*”) or via a four residue linker (dubbed “RC*4C*”) (Fig. 3a and Supplementary Table S2). Adapted RC proteins were expressed in *Rba. sphaeroides*. For LHCII, Lhcb apoproteins were expressed in *E. coli* and mature pigment-protein monomers refolded *in vitro* with purified pigments^43–46^. Three LHCII proteins were designed (Fig. 3b; see Supplementary Fig. S5a for protein sequences). The first, dubbed “dLHCII”, lacked twelve dispensable N-terminal amino acids that are not resolved in available X-ray crystal structures^30–32^ and had a His-tag at its C-terminus. The remaining two had either a truncated SpyTag variant (SpyTagΔ) added to the N-terminus of the truncated Lhcb1 (termed Td-dLHCII) or the full SpyTag sequence added to the C-terminus of the full Lhcb1 (termed LHCII-T) (Fig. 3b).

**Fig. 3.**
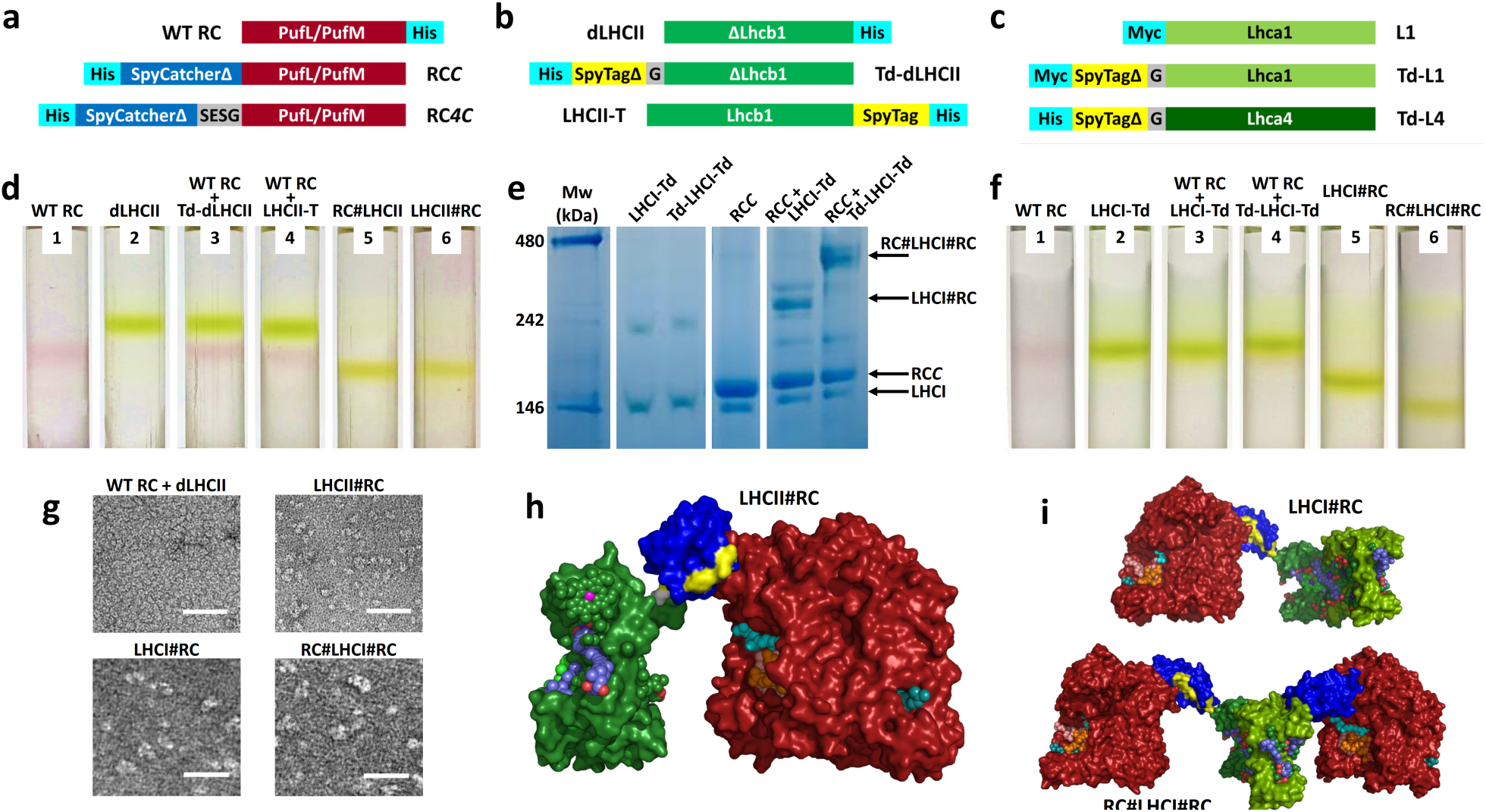
Engineering and assembly of RC-LHC chimeras. **a** Construct designs for adaptation of the RC. For purification the WT RC was modified with a His-tag on PufM. **b** Construct designs for adaptation of LHCII. The control LHCII was truncated at its N-terminus (dLHCII – see text) and was His-tagged at its C-terminus. **c** Construct designs for adaptation of LHCI which is a Lhca1/Lhca4 heterodimer. For **b** and **c** protein sequences are given in Supplementary Fig. S5a. **d** Sucrose density gradient fractionation of RCs (red bands) and LHCIIs (green bands). RC-LHCII chimeras migrate to a lower position in gradients than either RC or LHCII monomers, with no dissociation into components. **e** Blue native PAGE showing the formation of high molecular weight products by mixing LHCI-Td or Td-LHCI-Td with RC*C* (see Supplementary Fig. S7a for the full gel with more combinations). The multiple bands seen for the high molecular weight products are likely to be due to conformational heterogeneity. **f** Sucrose density gradient fractionation of RCs (red bands) and LHCIs (green bands). LHCI#RC chimeras and larger RC#LHCI#RC chimeras migrate to lower positions than either RCs or LHCI. **g** TEM images of an equimolar mixture of the WT RC and dLHCII (top/left), the LHCII#RC chimera (top/right), the LHCI#RC chimera (bottom/left) and the RC#LHCI#RC chimera (bottom/right). Additional images shown in Supplementary Fig. S17. **h** Molecular model of the LHCII#RC chimera. The RC (maroon) N-terminally adapted with SpyCatcherΔ (blue) is covalently linked to LHCII (green) C-terminally adapted with SpyTag (yellow). Cofactor colours are as described in Supplementary Fig. S1. **i** Molecular models of the LHCI#RC and RC#LHCI#RC chimeras. Colours as for panel **h** and Supplementary Fig. S1.

Adapted heterodimeric LHCI proteins (Fig. 3c; see Supplementary Fig. S5a for protein sequences) were also refolded from apoproteins expressed in *E. coli* ^34,38,47,48^. This involved mixing SpyTagΔ-adapted Lhca4 protein (Td-L4) with either unadapted Lhca1 protein (L1) or SpyTagΔ-adapted Lhca1 protein (Td-L1), to produce LHCI either singly or doubly modified with SpyTagΔ (termed LHCI-Td and Td-LHCI-Td, respectively). This enabled the creation of chimeras between LHCI and either one or two RCs.

### Self-assembly of RC-LHC chimeras

Following ultracentrifugation, purified RCs and LHCIIs could be visualised on sucrose density gradients as either a red or green band, respectively (Fig. 3d, gradients 1,2), and these two proteins also migrated separately in gradients loaded with a mixture with only either SpyTag or SpyCatcher adaptations (Fig. 3d, gradients 3,4). In contrast, mixing any SpyCatcherΔ-adapted RC with any SpyTag(Δ)-adapted LHCII produced a product, dubbed a “chimera”, that migrated further than either monomeric protein. The two examples shown in Fig. 3d (gradients 5,6) are chimeras from a RC*4C*/Td-dLHCII mix (dubbed “RC#LHCII”) and from a RC*C*/LHCII-T mix (dubbed “LHCII#RC”). The symbol “#” denotes the spontaneously-formed SpyCatcher/SpyTag interface domain. Chimera formation could also be detected on a native blue gel (Supplementary Fig. S6a). SDS-PAGE combined with western blotting using anti-His antibodies confirmed that chimera self-assembly was due to the formation of a covalent bond between the SpyTag(Δ)-adapted Lhcb1 polypeptide of LHCII and the SpyCatcherΔ-adapted PufL polypeptide of the RC (Supplementary Fig. S6b). The reaction half-time for chimera formation varied between 10 and 90 mins depending on the particular combination of adapted RC and LHCII (detailed in Supplementary Section 1).

LHCI-RC chimeras could also be assembled by incubation of LHCI-Td or Td-LHCI-Td with a three-fold excess of RC*C*. This again produced higher molecular weight products that could be separated from unreacted RCs on blue native gels (Fig. 3e). As designed, assembly of RC*C* with doubly-adapted Td-LHCI-Td complexes produced higher molecular weight products than with singly-adapted LHCI-Td complexes (Fig. 3e, right). Equivalent results were obtained with LHCI adapted with the full SpyTag and also with RC*4C* (Supplementary Fig. S7a). Analysis by SDS-PAGE and western blotting showed that chimera self-assembly was due to formation of a covalent bond between the SpyCatcherΔ-adapted PufL of the RC and Lhca4 of a singly SpyTagΔ-adapted LHCI (to form chimera LHCI#RC) or Lhca4 and Lhca1 of a doubly SpyTagΔ-adapted LHCI (to form chimera RC#LHCI#RC) (Supplementary Fig. S7b). Sucrose density gradient ultracentrifugation (Fig. 3f) showed that LHCI#RC chimeras (gradient 5) were clearly larger than LHCI alone (gradients 2-4) or unadapted RCs (gradients 1,3,4), and RC#LHCI#RC chimeras (gradient 6) were larger again.

Covalent-locking of the structure enabled purification of all LHCI-RC and LHCII-RC chimeras by a combination of nickel affinity and size-exclusion chromatography, absorbance spectroscopy being used to identify fractions containing protein oligomers with the designed molar ratio (Supplementary Fig. S8).

A change in protein morphology on chimera formation could be observed by transmission electron microscopy (TEM). Images of a mix of unadapted WT RCs and dLHCII showed a large number of monodispersed, regularly-sized objects of <10 nm diameter (Fig. 3g, top/left), whereas images of the purified LHCII#RC chimera revealed two-domain objects (Fig. 3g, top/right). The purified LHCI#RC and RC#LHCI#RC chimeras presented as objects with a more elongated morphology owing to the presence of one or two RCs and the heterodimeric LHCI (Fig. 3g, bottom). Molecular models of these chimeras, based on available X-ray crystal structures for the RC, LHCII, LHCI and SpyCatcher/Tag, are shown in Fig. 3h and 3i.

### Chlorophyll to bacteriochlorophyll energy transfer in chimeras

In solution, LHCII emission was quenched within each chimera in comparison to a control sample formed from an equivalent mix of the SpyTag-adapted LHCII and WT RCs (Fig. 4a; see spectra and other combinations in Supplementary Fig. S9). This was indicative of energy transfer, likely through a FRET mechanism at the distances implied by the chimera models (Fig. 3h,i), that was activated in these proteins in dilute solution by physically-linking the RC to the LHCII. These trends, observed with 650 nm excitation, were also seen in data on the same complexes obtained with other three excitation wavelengths, with no variation in emission spectrum line-shape (Supplementary Fig. S9). As well as being diagnostic of correctly refolded LHCII proteins, this lack of dependence of emission spectrum on excitation wavelength showed that the reduction in LHCII emission on chimera formation was not due to parasitic RC absorbance, which would be expected to be wavelength dependent (and also seen when WT RCs were mixed with each LHCII).

**Fig. 4.**
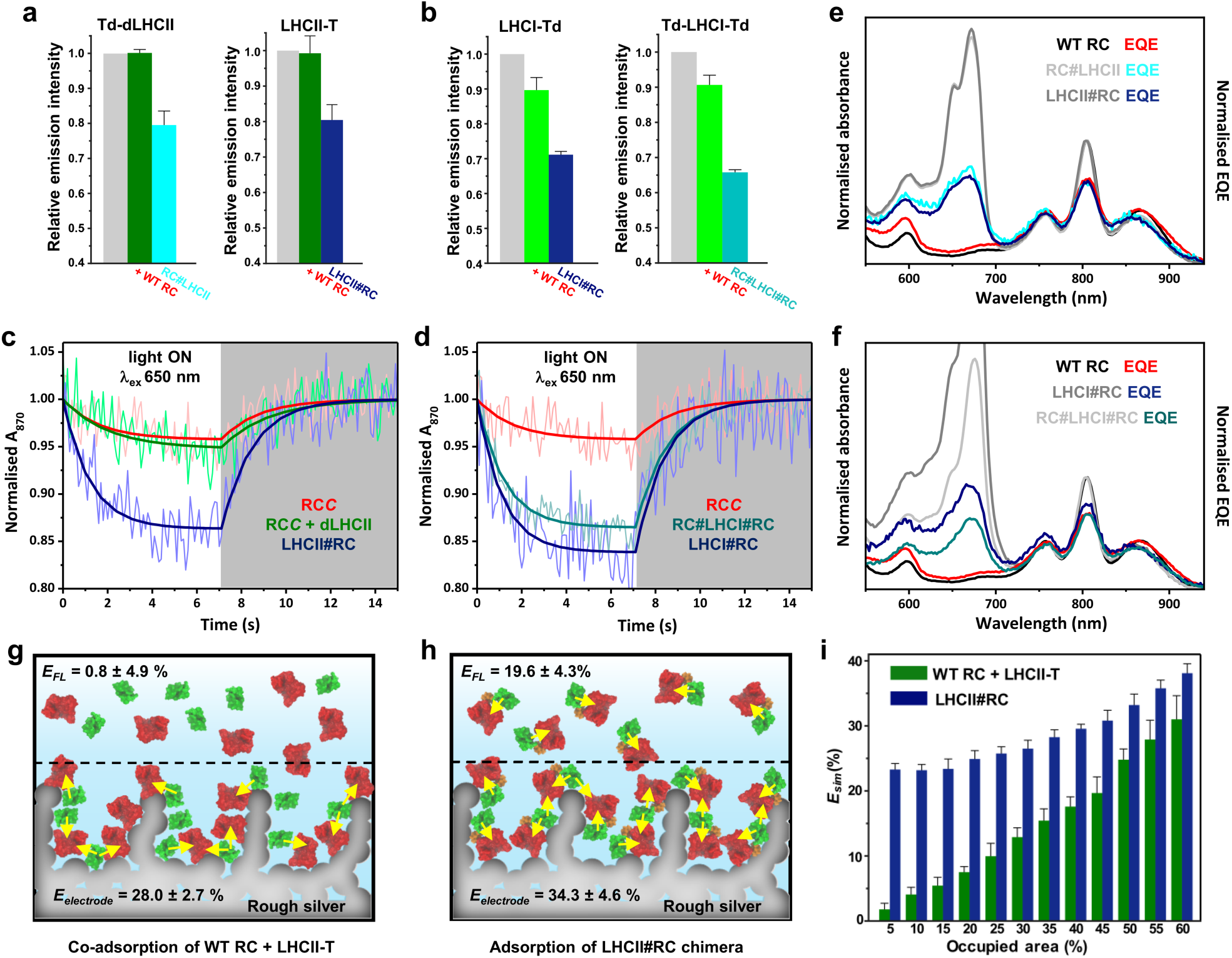
Energy transfer in chimeras in solution and on surfaces. **a** Emission at 682 nm from (left) the Td-dLHCII protein alone (grey), after addition of a two-fold excess of WT RCs (green) and in a LHCII#RC chimera (cyan) formed on mixing with a two-fold excess of RC*C*, and (right) equivalent data for LHCII-T. **b** Emission from (left) LHCI-Td alone (grey), after addition of a three-fold excess of WT RCs (green) and in a LHCI#RC chimera (navy) formed on mixing with a three-fold excess of RC*C*, and (right) equivalent data for Td-LHCI-Td. **c** Data and fits for photobleaching and dark recovery of P870 absorbance in RC*C*, a 1:1 RCC plus dLHCII mixture and the LHCII#RC chimera. **d** Data and fits for photobleaching and dark recovery of P870 absorbance in RC*C* and the two RC-LHCI chimeras. **e** Solution absorbance and EQE spectra for WT RCs compared with those for the two LHCII-RC chimeras. **f** Solution absorbance and EQE spectra for WT RCs compared with those for the two RC-LHCI chimeras. **g** Schematic of adsorption of independent RC (red) and LHCII (green) complexes on an electrode. Yellow arrows indicate possible energy transfer connections. **h** Equivalent schematic of adsorption of RC#LHCII chimeras. **i** Simulated apparent ET efficiencies as a function of packing density for independent RC and LHCII proteins or the LHCII#RC chimera.

To determine the fate of transferred energy, measurements of RC P870 photo-oxidation in response to 650 nm excitation were carried out on the LHCII#RC and RC#LHCII chimeras and fitted to a simple interconversion reaction (Eq. 1; all parameters are summarized in Supplementary Table S3). Bleaching of 870 nm absorbance was stronger in LHCII#RC chimeras than in controls comprising the RC*C* protein alone or a mixture of RC*C* with unadapted dLHCII complexes (Fig. 4c). The same was found for the RC#LHCII chimera (Supplementary Fig. S10a). Hence, decreased emission by the LHCII energy donor was accompanied by enhanced photo-oxidation of the RC energy acceptor, confirming energy transfer between the two proteins in solution that was switched on only after linking them by the SpyCatcher/Tag domain.

Turning to LHCI, a greater reduction of LHCI emission was seen on forming either LHCI#RC or RC#LHCI#RC chimeras than after mixing the same adapted LHCI proteins with WT RCs (Fig. 4b; and other combinations are shown in Supplementary Fig. S11). This effect was again seen to be independent of excitation wavelength (Supplementary Fig. S11a) showing it was not due to the absorbance of excitation light by the tethered RC(s). This emission quenching was accompanied by significant enhancement of P870 photo-oxidation in LHCI chimeras with one or two RC*C*, compared to that seen with RC*C* alone (Fig. 4d), confirming energy transfer. Doubly modified RC#LHCI#RC complexes showed less P870 bleaching than LHCI#RC complexes due to two tethered RCs competing for the exciton reservoir rather than one (see Supplementary Section 4 for more detail on the double-acceptor/single-donor FRET scheme).

Purified chimeras were also adhered to nanostructured silver electrodes to test their functionality. All were able to generate photocurrents, showing that dynamic interactions between the RC, cyt *c* and ubiquinone at the electrode surface, required for the generation of a photocurrent, were not obstructed by attaching the RC to LHCII or LHCI. All EQE action spectra recorded for chimeras exhibited low energy (Fig. 4e,f) and high energy chlorophyll bands (Supplementary Fig. S4c,d,g,h, green shading) indicating photocurrent generation powered by LHC absorbance.

### Energy transfer efficiency in chimeras

Apparent efficiencies of energy transfer from LHCII or LHCI to the RC in solution were estimated either from data on emission of the LHC energy donor (*E*_*FL*_) or from data on photo-bleaching of the RC energy acceptor (*E*_*P870*_) (see Materials and Methods, Eqs. 2-4). Efficiency *E*_*FL*_ was based on the additional quenching of LHC emission in a chimera relative to that in a compositionally-matched mixture of the relevant LHC variant and WT RCs (Eq. 2,3) or additional quenching in a LHC/WT RC mixture relative to that in a concentration-matched LHC-only sample. Efficiency *E*_*P870*_ was based on the enhanced rate of RC P870 photobleaching in a chimera relative to a matched RC-only control (Eq. 4).

Values of *E*_*P870*_ calculated from experimental data are shown in Table 1. The efficiency of energy transfer was low in mixtures of WT RCs with SpyTag adapted LHCIIs or LHCIs, consistent with expectations for a dilute (500 nM) solution of two proteins with no propensity to associate (see Supplementary Fig. S12 and Supplementary Table S4 for other control combinations). In marked contrast, *E*_*P870*_ was over 20 % in the corresponding RC-LHCII or RC-LHCI chimera (Table 1). For all chimeras the value of *E*_*FL*_ derived from LHC emission data was in excellent agreement with the values of *E*_P870_ derived from RC absorbance data (Table 1). This correspondence between independently-determined efficiencies from separate data sets reinforced the conclusion that energy transfer was taking place from the plant LHCs to the bacterial RCs within the chimera.

**Table 1.** Apparent energy transfer efficiencies and associated parameters.

| System | $E_{P870}$<br>(%) | $E_{FL}$<br>(%) | $E_{electrode}$<br>(%) | $U_{LHC}$<br>(%) |
| --- | --- | --- | --- | --- |
| LHCII-T + WT RC | $1.2 \pm 1.4^a$ | $0.8 \pm 4.9$ | $28.0 \pm 2.7$ | 106.7 |
| LHCII#RC | $22.8 \pm 5.1$ | $19.6 \pm 4.3$ | $34.3 \pm 4.6$ | 164.5 |
| RC#LHCII | $21.0 \pm 5.5$ | $20.3 \pm 3.6$ | $33.6 \pm 5.5$ | 176.6 |
| LHCI-Td + WT RC | $1.3 \pm 1.3^{a,b}$ | $10.3 \pm 3.6^b$ | $16.7 \pm 1.9$ | 120.1 |
| LHCI#RC | $20.2 \pm 4.6$ | $20.7 \pm 0.9$ | $24.8 \pm 1.6$ | 222.9 |
| WT RC + Td-LHCI-Td | $3.3 \pm 2.1^{a,b}$ | $9.3 \pm 2.8^b$ | $34.5 \pm 2.3$ | 129.8 |
| RC#LHCI#RC | $29.1 \pm 6.3^c$ | $27.4 \pm 1.8^c$ | $29.0 \pm 1.3$ | 90.9 |

<sup>a</sup> These low apparent energy transfer efficiencies may have arisen from some reabsorption of LHC fluorescence by RCs or a small degree of aggregation. Data for additional control mixtures can be found in Supplementary Table S4.
<sup>b</sup> The variance between $E_{P870}$ and $E_{FL}$ in these two cases is attributed to the latter largely reflecting a decrease in LHCI quantum yield on adding WT RCs rather than being due to ET (see text).
<sup>c</sup> RCs conjugated to each of Lhca1 and Lhca4 in the LHCI heterodimer.

“On electrode” apparent energy transfer efficiencies (*E*_*electrode*_) were also determined from the EQE action spectra, as described in Materials and Methods. In general, values of *E*_*electrode*_ were higher than either estimate of energy transfer efficiency in solution (Table 1). This was particularly striking for mixtures of WT RCs and SpyTag(Δ)-adapted LHCII or LHCI (shown schematically in Fig. 4g) where energy transfer in solution had a very low apparent efficiency. However, for the RC/LHCII chimeras in particular the value of *E*_*electrode*_ was also substantially higher than *E*_*P870*_ or *E*_*FL*_ (Table 1 and Fig. 4h), suggesting that adhering the chimeras to a surface turned on inter-chimera ET that supplemented the intra-chimera ET also observed in solution. This effect was less pronounced for the RC/LHCI chimeras, particularly for complex RC#LHCI#RC where there were already two RCs per LHCI antenna (Table 1).

To examine whether the benefits of pre-linking RCs and LHCs in a chimera would be enhanced across a range of packing densities, a 2D Monte Carlo simulation was carried out as detailed in Supplementary Sections 2 and 3 (and summarized in Supplementary Fig. S13). In this either LHCII#RC chimeras or a mixture of LHCII-T and WT RC proteins were represented as hard-discs on a 2D surface and centre-to-centre distances calculated as a function of packing density. The outcome of this simulation was an apparent energy transfer efficiency (*E*_*sim*_) based on how protein packing densities affected mean inter-protein distances (Fig. 4i). In the high packing regime, *E*_*sim*_ was in good agreement with the slightly higher *E*_*electrode*_ determined from chimeras than a mixture of unadapted proteins. As the packing density dropped to a low value (right to left in Fig. 4i), *E*_*sim*_ for the chimeras gradually declined to around the estimates of *E*_*P870*_ and *E*_*FL*_ for the LHCII#RC chimera in solution (22.8%/19.6%). In contrast, *E*_*sim*_ for the protein mixture declined steeply to less than 2% at the lowest packing density, again in agreement with estimates of *E*_*P870*_ and *E*_*FL*_ for the protein mixture in solution (1.2%/0.8%). This reinforced the conclusion that pre-tethering of the RC and LHCII protein into a chimera brought an added benefit even under conditions where co-localisation of the proteins on a surface switched on energy transfer between the two irrespective of tethering. Presumably pre-tethering can mitigate against situations where, for example, formation of RC-rich or LHCII-rich sub-domains and sub-optimal mixing can lead to some proteins being outside the FRET distance (Supplementary Fig. S14, marked with blue triangles).

The EQE spectra were also used to estimate the percentage improvement in the use of visible light by an LHC/RC bio-photoelectrode compared to a RC bio-photoelectrode. Consistent with values of *E*_*electrode*_, the presence of an LHC consistently boosted the use of visible light, with the strongest effects seen for electrodes fabricated from chimeras.

## Discussion

The data establish that it is possible to genetically encode *in vitro* self-assembly of a hybrid chlorophyll/bacteriochlorophyll solar energy conversion system using a highly-specific split-interface domain. To our knowledge such combinations of chlorin and bacteriochlorin pigments are not used for light harvesting in nature, although in green sulphur bacteria the multiple BChl *a* light harvesting and electron transfer cofactors of the RC are supplemented by four molecules of Chl *a* that are used electron acceptors during charge separation^49^. In a similar vein, in the related heliobacterial RC the multiple BChl *g* (an isomer of Chl *a*) cofactors are supplemented by two molecules of 8^1^-hydroxychlorophyll *a* that also act as electron transfer acceptors^50^. Hence some organisms have evolved to supplement bacteriochlorin cofactors with chlorins to achieve charge separation, but not to expand solar energy harvesting in the way demonstrated here.

The SpyCatcher/Tag system provided a versatile means of constructing self-assembling hybrid photosystems. LHCII could be modified with SpyTag at either its N- or C-terminus, and by also using heterodimeric LHCI proteins that were either singly or doubly SpyTag modified the oligomeric state of the chimeras could be varied between heterodimers (RC#LHCII and LHCII#RC), heterotrimers (LHCI#RC) and heterotetramers (RC#LHCI#RC). The SpyCatcher/Tag linking domain produced predictable and stable products due to its very high partner specificity and the autocatalytic formation of a locking covalent bond. This binding reaction, which under the present conditions was found to have a half time of between 10 and 90 minutes, was irreversible, relatively insensitive to reaction conditions and was free of side products (i.e. a failed reaction did not lead to depletion of reactants). The assembly strategy used, using *E. coli* and *Rba. sphaeroides* as separate bacterial factories for the synthesis of protein components that could be assembled *in vitro*, avoided the need to re-engineer a host organism to be able to produce both chlorophyll and bacteriochlorophyll (and different types of carotenoid). This methodology therefore provides a route for the bottom-up redesign of a photosystem *in vitro* despite the challenges of working with large, multi-component integral membrane complexes.

The mechanism of solar energy conversion operating in the chimeras, based on the well-understood photophysical properties of the component proteins, is summarised in Fig. 5. Energy captured by the pigment systems of LHCII or LHCI will be passed to the RC in a downhill manner, exciting the primary electron donor bacteriochlorophylls (P870*) and initiating charge separation to form P870^+^Q_B_^-^. Energy harvested by the chlorophyll *b* (or carotenoid – not shown) pigments of either LHC is passed to the lower energy chlorophyll *a*. Inter-protein energy transfer is likely to involve a sub-set of red-shifted chlorophyll *a* in either LHC and entry of energy into the RC is likely to occur principally via the bacteriopheophytin cofactors (H_A/B_) as their absorbance has the greatest spectral overlap with LHC emission (Fig. 1b,c).

**Fig. 5.**
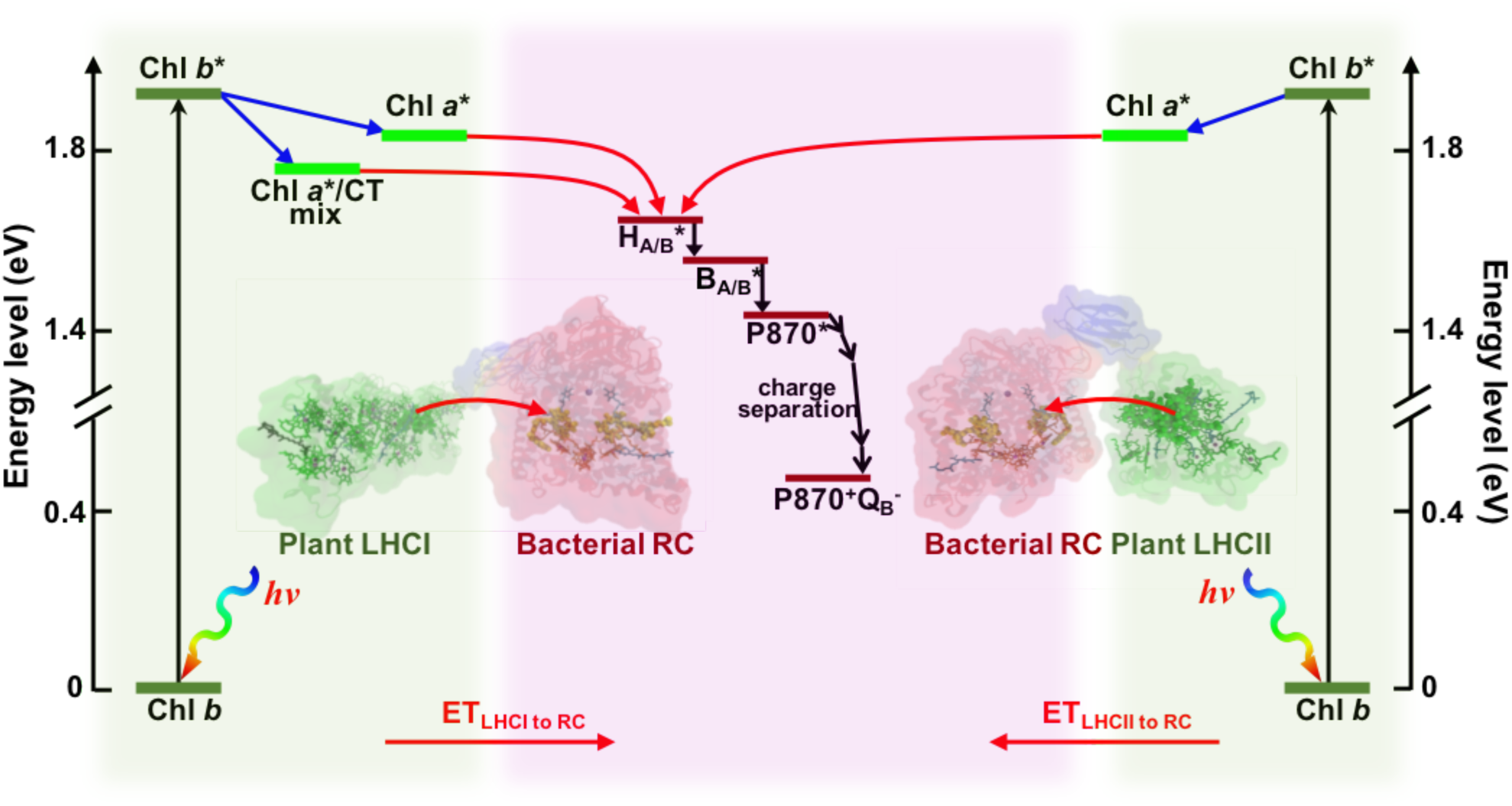
Solar energy conversion in chimeras. Energy flow within LHCII or LHCI is from higher energy chlorophyll *b* to lower energy chlorophyll *a*. LHCI also exhibits a red-shifted emissive state with mixed excitonic/charge transfer (CT) character. Excited state energy entering the RC via the bacteriopheophytins (H_A/B_) migrates to the P870 bacteriochlorophylls via the monomeric bacteriochlorophylls (B_A/B_), initiating charge separation to form P870^+^Q_B_^-^. Energy harvested by the carotenoid pigments of LHCII or LHCI (not shown) would transferred to the RC via their chlorophylls through fast internal relaxation^61^.

As evident from comparing Fig. 1c with Fig. 1b, LHCI exhibits a red-enhanced fluorescence that produces a ∼80% stronger spectral overlap with RC absorbance (factor *J* in Supplementary Table S1) compared to LHCII. Despite this, the efficiency of ET in the LHCI#RC chimera was not significantly higher than that in either the LHCII#RC or RC#LHCII chimera. This is likely due to the reconstituted LHCI heterodimers being in a partially quenched state^48,51^ that reportedly reduces their quantum yield to only 29 % of that of LHCII^36^, so counteracting the potential benefits of an enhanced spectral overlap. In agreement with this our estimates of quantum yield were 30% for LHCI-Td and 28% for Td-LHCI-Td (Supplementary Table S1). In future work it might be possible to partially overcome this through SpyTag modification of LHCI in a native organism, as the quantum yield of purified native LHCI has been reported to be ∼64% that of LHCII, more than double that of recombinant LHCI^36^.

Estimates of ET efficiency in RC#LHCI#RC chimeras in solution were consistently higher than those for the LHCI#RC chimera (parameters *E*_*P870*_ and *E*_*FL*_ in Table 1), consistent with the presence of two ET acceptors in the former. Estimates of the ET efficiency to the second RC added to Lhca1 in RC#LHCI#RC, made using Eq. 5, yielded values that were either 50 % or 69 % of that for transfer to the first RC attached to Lhca4. This is consistent with the presence of a relatively low energy red-form chlorophyll *a* dimer in the Lhca4 subunit (Supplementary Fig. S1d) that is responsible for the red-enhancement of the LHCI emission spectrum^36,38,47,48^, and which may have produced more efficient ET to the RC attached to Lhca4 than that attached to Lhca1.

## Conclusions

This work shows that genetically adapting two diverse photosynthetic membrane proteins with the components of an extramembrane interface domain enables *in vitro* self-assembly of a chimeric photosystem in which UV/near-IR solar energy conversion by a bacteriochlorophyll-based RC is augmented by visible light capture by chlorophyll-based LHCs. This approach inspired by a concept of synthetic biology, to adapt naturally incompatible biological modules to interface in a standardized way through genetic encoding, creates covalently stabilised macromolecular photosystems that are predictable and programmable. In addition to providing novel photosynthetic structures and energy transfer pathways to explore, these polychromatic photosystems constitute interesting new materials for biohybrid devices that in recent years have expanded in application beyond photoelectrochemical solar energy conversion to fuel molecule synthesis, energy storage, biosensing, touch sensing and photodetection. Finally, the demonstrated flexibility with which RCs and LHCs could be interfaced opens the possibility of constructing more elaborate, self-assembling chimeric photosystems that employ multiple orthogonal linking modules^52,53^ and a wider range of photosynthetic and redox proteins that, despite being separated by billions of years of evolution, can be adapted for future solar energy conversion through genetic programming of standardized interfaces.

## Acknowledgements

The Lhcb1.3 plasmid was a kind gift from Prof. Roberta Croce of the Vrije Universiteit Amsterdam, The Netherlands. We also thank Dr. Majid Mosayebi from the School of Mathematics, University of Bristol for generous advice on simulations. J.L and M.R.J acknowledge funding from the EPSRC/BBSRC Synthetic Biology Centre for Doctoral Training (EP/L016494/1) and from the BrisSynBio Synthetic Biology Research Centre at the University of Bristol (BB/L01386X/1). R.N.F acknowledges support from the Netherlands Organisation for Scientific Research (NWO) for a Vidi grant, and V.M.F for funding from NWO Veni project 16866.

## Author contributions

J.L and M.R.J conceived the research. J.L engineered the chimeras, characterised binding and energy transfer and conducted the simulations. V.M.F and J.L carried out photochronoamperometry under supervision from R.N.F. J.L and M.R.J drafted the manuscript and all authors commented on the manuscript. M.R.J supervised the project.

## Competing interests

The authors declare no competing interests.

## Methods

### RC modification and purification

SpyCatcherΔ, a version of the SpyCatcher protein lacking nine C-terminal amino acids that are not resolved in the X-ray crystal structure of SpyCatcher/SpyTag^42^ was used in order to reduce the length of the linking peptide between the main bodies of the RC and SpyCatcher proteins. It was fused to the N-terminus of the RC PufL polypeptide, which is an alanine exposed at the protein surface on the cytoplasmic side of the membrane (white spheres in Supplementary Fig. S1a), either directly or via a four (SESG) amino acid linker, and was preceded by a His-tag for purification (Fig. 3a). Nomenclature used for these and all other component proteins and chimeras are summarised in Supplementary Table S2. The required plasmids were constructed using synthetic DNA (Eurofins). The WT RC control was also modified with a His tag for purification as described previously^54^. All RCs were expressed in a strain of *Rba. sphaeroides* engineered to lack light harvesting complexes^54,55^, were purified by nickel affinity chromatography and gel filtration^54^, and were stored at −80 °C as a concentrated solution in 20 mM Tris (pH 8.0)/0.04 % DDM (Tris/DDM buffer).

### LHCII modification, refolding and purification

To enable low-cost and rapid genetic modification of plant LHCII the designed apoproteins were expressed in *E. coli* and the mature pigment-proteins refolded in DDM using pigments purified from spinach according to a well-established protocol^43^. Designs of the modified LHCII are outlined in Fig. 3b and Supplementary Fig. S5a. Complex dLHCII lacked twelve dispensable N-terminal amino acids (Supplementary Fig. S1d). This sequence, which includes four basic amino acids, is involved in stacking of thylakoid grana but can be removed without affecting core LHCII light-harvesting function^31^. Their removal minimised the sequence linking the main body of LHCII (starting at serine 14) to additional components added at the N-terminus. In the second construct a truncated SpyTagΔ was added to the N-terminus of dLHCII to form Td-dLHCII. This modified SpyTagΔ lacked three dispensable amino acids at its C-terminus^42^, further reducing the linker to the N-terminus of LHCII. In the third construct the full 13 amino acid SpyTag peptide was added to the C-terminus of Lhcb1 after a three amino acid linker (termed LHCII-T). In all cases a His-tag placed adjacent to the SpyTag sequence ensured the latter was retained in the final, purified pigment-protein (Fig. 3b). A fourth LHCII with the full SpyTag on the N-terminus of the truncated Lhcb1 was also constructed (T-dLHCII) but gave identical results to Td-dLHCII (shown in Supplementary Fig. S5 and data shown in some other supplementary Figures).

The starting point for production of the designed LHCII holoprotein was a pET-28a expression vector containing a gene encoding the Lhcb1.3 protein from *Arabidopsis thaliana* (UniProtKB entry P04778), that was a kind gift from Prof. Roberta Croce, Vrije Universiteit Amsterdam. Modification of this gene was carried out by Gibson assembly using oligonucleotides sourced from Eurofins or using the Q5^®^ Site-Directed Mutagenesis Kit from NEB. Designed apoproteins were expressed in *E. coli* Rosetta™ 2 (Novagen), purified from inclusion bodies, and the mature pigment-protein assembled *in vitro* using purified chlorophyll *a*, chlorophyll *b* and carotenoid pigments with DDM as the supporting detergent, according to a previously published protocol^43^ with exception that ß-mercaptoethanol was replaced by dithiothreitol (DTT). Pigments were purified as described previously^43^ with the exception that they were dried from acetone solution using a freeze drier rather than a centrifugal vacuum concentrator. Pigment concentration and composition was determined in 80 % (v/v) acetone using published equations^56^.

Each refolded LHCII was first purified by nickel affinity chromatography and then by gel filtration chromatography. Fractions with the lowest A_470_ to A_674_ ratio and invariant emission profiles in response to 440 nm, 475 nm and 500 nm excitation were kept and pooled. Purified proteins were stored at −80°C before use as concentrated solutions in Tris/DDM. An extinction coefficient at the chlorophyll *a* Q_y_ band of 546,000 M^-1^ cm^-1^ was used to estimate LHCII concentration^25^.

The refolded LHCII complexes had absorbance spectra that were similar to one another (Supplementary Fig. S5b) and to spectra previously published by others^43–46^. Their emission spectra were highly similar (Supplementary Fig. S5c), and the line-shapes of these spectra were invariant with excitation wavelength (Supplementary Fig. S5d), a feature diagnostic of a structurally-intact LHCII. Pigment compositions were similar to those typically reported for recombinant LHCII (Supplementary Fig. S5e)^46^.

### LHCI modification, refolding and purification

LHCI heterodimeric complexes were assembled from modified versions of *A. thaliana* proteins Lhca1 (UniProtKB entry Q01667) and Lhca4 (UniProtKB entry P27521). The mature Lhca1 was modified at its N-terminus either with a Myc protein purification affinity tag (termed L1) or with a Myc-tag followed by the shortened ten amino acid SpyTagΔ (termed Td-L1), and the mature Lhca4 was modified at its N-terminus with a His-tag followed by SpyTagΔ (termed Td-L4) (Fig. 3c and Supplementary Figure S5a). combining L1 or Td-L1 with Td-L4 produced LHCI heterodimers with either one (LHCI-Td) or two (Td-LHCI-Td) SpyTag adaptations.

Expression plasmids were pET-28a containing synthetic genes sourced from Eurofins. Following apoprotein expression in *E. coli*, LHCI heterodimers were assembled by refolding with purified pigments in octyl glucoside (OG)^34,36,38,47,48^. A 20 % excess (by mass) of either L1 or Td-L1 was mixed with Td-L4 to reduce the level of free Td-L4 monomer after refolding. The apoprotein:total pigment ratio was kept the same as for LHCII refolding. Nickel affinity chromatography was used to separate the His-tagged LHCI dimer from residual Lhca1 monomer (which was not His-tagged). Each LHCI was then further purified by gel filtration chromatography and stored at −80°C before use as a concentrated solution in Tris/DDM. An extinction coefficient for the chlorophyll *a* Q_y_ band equal to 1,092,000 M^-1^ cm^-1^ was used to evaluate LHCI concentration since its chlorophyll *a* content is approximately twice that of a refolded LHCII monomer^34^.

### Chimera formation and verification

The standard approach to chimera formation was to mix RCs with a two-fold molar excess of LHCII, or a three-fold molar excess of LHCI, and then separate the chimera from unreacted components by gel filtration chromatography, using absorbance spectroscopy and each constituent molar extinction coefficient to assess the RC:LHC molar ratio in each column fraction (see Supplementary Fig. S8).

Formation of chimeras was initially verified by sucrose density gradient ultracentrifugation (Fig. 3d,f). Linear sucrose gradients were prepared by freezing and thawing 10 mL of 21 % (w/v) sucrose in 20 mM Tris/0.04% DDM (pH 8.0). Each gradient was loaded with 400 μL of sample with each photoprotein at a concentration of 2.5 μM and then capped with 1 mL of 20 mM Tris/0.04% DDM (pH 8.0). Gradients were ultracentrifuged in a Sorvall TH-641 swing-out rotor at 38,000 rpm for 18 hours at 4 °C.

The protocol for blue-native PAGE was adapted from one published previously^57^. Precast NativePAGE 4-20 % gels purchased from Thermo were run in a Bis-Tris buffer system. Coomassie blue dye at 0.02 % (w/v) was used in the cathode buffer but not in the loading buffer. The gel cassette was placed in an ice bath and run at 150 V for 1 h followed by 250 V for 2 h.

SDS-PAGE was carried out using precast 4-20 % gradient gels (Bio-Rad). A standard loading of 20 pmol RC was used. Loaded gels were run at 200 V for 45 mins and stained overnight at room temperature with Quick Coomassie Stain (Generon™).

Western blotting was carried out following protein transfer onto a nitrocellulose membrane (GE Healthcare) on a TE 77 PWR Semi-Dry Transfer Unit (45 mA/gel and 30 min with a NOVA Blot kit) in 30 mins. The membrane was blocked overnight with 5 % milk PBS-Tween (PBS/T) buffer and then incubated with horse radish peroxidase (HPR) conjugated antibodies in the same buffer for 1 h. The membrane was developed using 1x LumniGLO(R) (CST®) after rinsing the membrane three times with PBS/T buffer. Finally, the result was recorded on an ODYSSEY imaging system (LI-COR Biosciences). Re-probing of the membrane was accomplished by stripping and a repeat process of incubation in 5 % milk PBS/T buffer. Stripping of membrane was achieved by incubating twice in mild stripping buffer (200 mM glycine, 0.1 % SDS (w/v), 1 % Tween 20 (v/v), pH 2.2) for 5 mins and twice in TBST buffer (50 mM Tris, 150 mM NaCl, 0.1 % Tween 20 (v/v), pH 7.6), before finally transferring in PBS/T.

### Spectroscopy

Absorbance spectra were recorded on a Varian Cary60 spectrophotometer and emission spectra on a Varian Cary Eclipse Fluorimeter in nitrogen-gassed, freshly prepared Tris/DDM.

Photo-oxidation of the RC P870 primary electron donor was measured using an optical fibre attachment for the Cary60 and a four-way cuvette holder (Ocean Optics, Inc.). For excitation, light from a HL-2000 light source (Ocean Optics, Inc.) was passed through an optical fibre and a 25 nm band-pass filter centred at 650 nm (Edmund Optics Ltd). Incident light intensity was approximately 0.1 mW cm^-2^, which excited ∼15 % of the RC population. Light-on/off was controlled using the electronic shutter on the light source triggered by a TGP110 pulse generator (Aim-TTi Ltd, UK). After incubation with a 10-fold excess of ubiquinone-0 (UQ_0_) in the dark for 10 mins, samples at a RC concentration of 0.5 μM (0.25 μM with LHCI-Td) were housed in a 3 mm path length, four-sided micro cuvette (110-15-QS, Hellma^®^ Analytics). Each measurement was repeated five times and the traces were fitted to a model assuming a simple interconversion between the ground and photo-oxidised state:

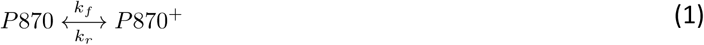

Parameters *k*_*f*_ and *k*_*r*_ from these fits are shown in Supplementary Table S2. All control samples had equimolar LHC and RC except a WT RC/Td-LHCI-Td mix where the molar ratio of RC to LHC was two.

### Quantum yield estimation

The quantum yield of LHCs was determined by comparison to the dye DCM (4-(dicyanomethylene)-2-methyl-6-(4-dimethylaminostyryl)-4H-pyran; Sigma) dissolved in methanol. To avoid self-shading the absorbance of LHCs and DCM was set around 0.07 across the relevant spectral region (Supplementary Fig. S15a). Emission from DCM and LHCII_H_ (average of 10 measurements; Supplementary Figure S15b) was corrected for spectral response and used to calculate their relative integral photon fluxes^58,59^. The value for Φ_*D*_ was estimated with reference to Φ_*DCM*_ = 0.435 ^60^ and the refractive indices of water (*n*_*water*_ = 1.333) and methanol (*n*_*methanol*_ = 1.328).

### Photochronoamperometry and EQE action spectra

Nanostructured silver electrodes of 2 mm diameter were prepared as described previously^20^. Pigment-proteins at concentrations between 20 µM and 100 µM were drop-casted onto prepared electrodes in the dark at 4 °C for one hour and unbound protein was removed by repeated mechanically-controlled dipping in 20 mM Tris (pH 8) at 4 °C. Coated electrodes were immersed in 20 mM Tris (pH 8)/50 µM KCl/20 µM horse heart cyt *c*/1.5 mM ubiquinone-0 (Q_0_) in a room temperature electrochemical cell fitted with an Ag/AgCl/3M KCl reference electrode and a platinum counter electrode. Photocurrents were measured at a bias potential of −50 mV vs Ag/AgCl, controlled by a PGSTAT128N potentiostat (Metrohm Autolab). Illumination was supplied by 870 nm or 650 nm LED (Roithner Lasertechnik) with irradiances of 32 or 6.7 mW cm^-2^, respectively, at the electrode surface with about 50 nm FWHM (full width at half maximum) for both. EQE action spectra were recorded using a tungsten-halogen source passed through a monochromator (Supplementary Fig. S16a)^20^. All control samples had equimolar LHC and RC except a WT RC/Td-LHCI-Td mix where the molar ratio of RC to LHC was two.

### Transmission electron microscopy

Negative stain TEM was carried out as described previously^27^ on an equimolar mixture of 500 nM WT RCs and dLHCII, 500 nM LHCII#RC heterodimers, 100 nM LHCI#RC or 100 nM RC#LHCI#RC. Samples were stained with 3% uranyl acid (UA) and imaged with a FEI Tecnai 12 120kV BioTwin Spirit TEM.

### Estimation of energy transfer efficiency

Apparent efficiencies of ET were calculated from LHC emission spectra (*E*_*FL*_) using:

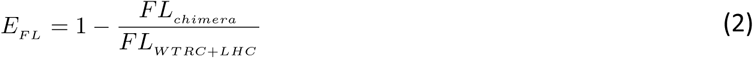

where *FL*_*chimera*_ was the intensity of LHC emission in a chimera and *FL*_*WTRC + LHC*_ was that in a concentration-matched mixture of the appropriate LHCII or LHCI variant and the WT RC. A similar approach was used for estimating the apparent ET efficiency in mixtures of WT RCs and LHCII or LHCI, expressing *FL*_*WTRC + LHC*_ as a function of the emission from the same concentration of the LHC (*FL*_*LHC*_). For LHCII, where the line shape of the emission spectrum did not vary as it is a single quantum system^61^, maximum emission values were used in Eq. 2 as a simple measure of emission intensity. For LHCI, which has multiple distinct emission states^38^, values of emission intensity (*FL*_*int*_) were produced by integration across the emission spectrum using Eq. 3, and then applied in Eq. 2.

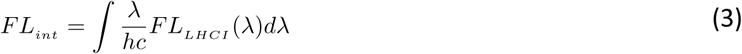

Apparent efficiencies of ET were also calculated from the rate of P870 photobleaching (*k*_*f*_) from the kinetic analyses summarised in Supplementary Table S2. To enable this the intensity of the 650 nm excitation light used in these experiments was kept low such that no more than ∼15 % of P870 was oxidised within the lifetime of P870^+^ (∼ 1s), ensuring that photooxidation directly represented the quantity of energy received by either direct absorption by the RC or ET from the tethered LHC. The apparent efficiency of ET (*E*_*P870*_) was estimated from the rate of P870 photobleaching using:

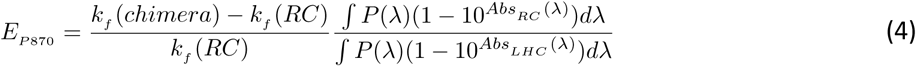

where *k*_*f*_ was the rate of P870 oxidation in a chimera (*chimera*) or the equivalent RC-only control (*RC*) (Supplementary Table S3). Integration of incident photon flux (*P*) and the 1-transmission of RCs or LHCs as a function of wavelength provided the number of photons absorbed by either RCs or LHCIIs per unit area per second (Supplementary Table S3).

Equation 4 was also used for estimation of *E*_*electrode*_, with parameter *k*_*f*_ replaced by the maximum EQE around 650 nm (i.e. the same illumination region as used for measurements of P870 oxidation). The efficiency was determined by comparing the LHC’s contribution to the EQE with the sample absorbance at the corresponding wavelength (Supplementary Fig. S4). In addition to enabling direct comparison of *E*_*electrode*_ and *E*_*P870*_, the data at 650 nm were not affected by parasitic absorption or unwanted emission from cyt *c*, UQ_0_ or nano-structured silver used in photocurrent measurements^20,40^.

For RC#LHCI#RC chimeras there were two acceptors per LHCI, one connected to the Lhca1 subunit and one to the Lhca4 subunit. Efficiencies of energy transfer to the Lhca1-connected RC (*E*_*a1*_) were estimated from:

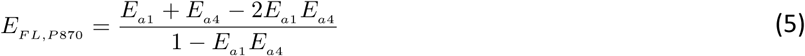

where *E*_*FL,P870*_ was the apparent energy transfer efficiency for the RC#LHCI#RC chimera estimated from either LHC fluorescence or P870 photobleaching and *E*_*a4*_ was the corresponding apparent energy transfer efficiency for the LHCI#RC chimera where the single RC is attached to Lhca4. From *E*_*FL*_ the value of *E*_*a1*_ was 10.4 % (compared to *E*_*a4*_ = 20.7 %) and from *E*_*P870*_ the value of *E*_*a1*_ was 13.9 % (compared to *E*_*a4*_ = 20.1 %).

### Estimation of solar radiance coverage enhancement

The effect of the LHCs on the performance of a bio-photoelectrode in response to visible light was estimated using:

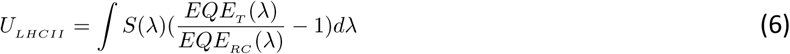

where *S(λ)* was the air mass 1.5 standard solar power reference spectrum as a function of wavelength (W m^-2^nm^-1^), *EQE*_*T*_ was the EQE spectrum of each LHC+RC system and *EQE*_*RC*_ was that of the RC-only reference (Supplementary Fig. S4, green versus red shade). Integration provided an estimate of the improvement in the use of solar energy (*U*_*LHCII*_) between 400 and 700 nm where the chlorophyll-based LHCs absorb. Values are compiled in Table 1.

### Protein structures and chimera modelling

Protein structures used in modelling were Protein Data Bank entries 3ZUW for the *Rba. sphaeroides* RC^62^, 2BHW for the LHCII from pea^31^, 4KX8 for the LHCI from pea^37^ and 4MLI for SpyCatcher/Tag^42^. Schematic models of chimeras were produced using Modeller^63^.

## Notes

#### Summary of Updates

Revised version (V26)

