## Supplemental Info for "Polychromatic solar energy conversion in pigment-protein chimeras that unite the two kingdoms of (bacterio)chlorophyll-based photosynthesis"

Supplementary Table S3. Summary of fitting of P870 photobleaching and computed photon absorbance. . 35

Supplementary Table S4 . Estimates of ET efficiency in additional control protein mixtures. ........ 36

***Supplementary References*** ........ ***36***

**Supplementary text**

**Section 1 – SDS-PAGE confirmed formation of RC-LHC fusion polypeptides mediated by SpyTag/SpyCatcher**

The WT RC used in this work had a His-tag on its PufM polypeptide and in SDS-PAGE (**Supplementary Fig. S6b**, lane 2) displayed three bands attributable to (bottom to top) PufL, $PufM_H$ and PuhA. For the two SpyCatcherΔ-modified RCs (**Supplementary Fig. S6b**, lanes 3,4) three bands were also seen corresponding to (bottom to top) unmodified PufM, PuhA and an enlarged PufL modified with SpyCatcherΔ and a His-tag. Although the theoretical molecular weight for this modified PufL is 44.4 KDa for RC*C* and 44.8 KDa for RC*4*C, it is well-known that due to the high hydrophobicity of PufL it does not run true to its molecular weight in SDS-PAGE. Despite this, the positioning of the bands for the SpyCatcherΔ-modified PufL proteins were consistent with PufL subunit being modified with the approximately 13 KDa SpyCatcher sequence (bottom band in lane 2 shifts to top band in lanes 3 and 4).

In addition to the two SpyTag(Δ)-modified LHCIIs described in the main text (Td-dLHCII and LHCII-T) a third LHCII with the full SpyTag at the N-terminus of dLHCII was also made (T-dLHCII). The approximate molecular weights of these proteins from amino acid sequences are 26.3, 27.9 and 26.7 KDa for Td-dLHCII, LHCII-T and T-dLHCII, respectively. Despite also being integral membrane proteins, these molecular weights correlated well with their positions in the SDS-PAGE analysis (**Supplementary Fig. S6b**, lanes 5-7).

On mixing each SpyTag(Δ)-modified LHCII with a two-fold excess of either SpyCatcherΔ-modified RC a new band appeared that ran between the 55 and 70 kDa markers (**Supplementary Fig. S6b**, lanes 8-13). The position of this isopeptide bond-locked product polypeptide correlated well with an expected molecular weight of approximately 64 KDa (i.e. a combination of a ~27 KDa SpyTag-LHCII and a SpyCatcher-modified RC PufL running at ~37 KDa). Moreover, as this LHCII-RC fusion peptide includes two His tags, its identity could be confirmed with an anti-His Western blot (**Supplementary Fig. S6b**, lower panel).

An equivalent analysis also confirmed the formation of either one or two fusion proteins when chimeras were formed between either singly or doubly SpyTag-modified LHCI and the two SpyCatcherΔ-modified RCs (**Supplementary Fig. S7b**). In this case a duplicate set of samples were analysed in which the full SpyTag sequence had been added either singly or doubly to LHCI (lane labels greyed-out in **Supplementary Fig. S7b**. Findings with this set of SpyTag-modified LHCI polypeptides were identical to those with the SpyTagΔ-modified polypeptides and are not mentioned in the main text.

**Section 2 – SDS-PAGE revealed the kinetics of chimera formation.**

SDS-PAGE could also be used to monitor the kinetics of chimera formation through emergence of the isopeptide bond-locked product polypeptide at higher molecular weight and the associated disappearance of SpyTag(Δ)-LHC and SpyCatcherΔ-RC bands at lower molecular weight (**Supplementary Fig. S18**). Western blotting with an anti-His antibody again verified the identities of these bands due to the presence of one or more His tags (**Supplementary Fig. S18**). **Supplementary Fig. S19** shows data for depletion of the SpyCatcher-PufL protein band, quantified by densitometry of the SDS-PAGE band, fitted with a bimolecular reaction scheme. This showed that chimera formation proceeded with a reaction half-time of between 11 and 91 mins in the six combinations tested. In comparison, it has been reported that the half-time for the reaction between a SpyTag-MBP (Maltose Binding Protein) construct and SpyCatcher is 2.4 mins.[1] The slower kinetics for the reaction between modified LHCs and RCs were likely due to the frustration caused by much larger adducts associated with the Spy-domains, and in support of this there was a general trend that the reaction rate became slower with increasing size of adduct (**Supplementary Fig. S19**, LHCII versus larger LHCI). In addition, the length of the linker joining the Spy-component with the adduct protein might also play a crucial role in determining the rate of chimera formation, as illustrated by the substantial difference between RCC and RC*4*C on mixing with LHCII-T (**Supplementary Fig. S19**, middle row). When the linker is short, a greater spatial constraint was applied to the reaction, the resulting large entropy barrier likely leading to the observed slow kinetics.

Despite these differences in kinetics, all chimeras could be purified from a completely reacted mixture after overnight incubation because of the irreversible nature of the reaction.

**Section 3 - Hard disc Monte Carlo simulation**

To explore the influence of protein packing density on ET efficiency a Monte Carlo simulation was carried out in which RC and LHCII proteins were represented as hard discs on a 2D surface with periodic boundary conditions (**Supplementary Fig. S13**). The X-ray crystal structures of the RC and LHCII proteins were used to prepare a 2D projection in the plane of the membrane with a 1.5 nm thick outline added to the projected shape to represent the DDM detergent micelle[2].

The simulation followed an athermal Metropolis Monte Carlo approach that considered the protein complexes as hard discs. First, discs representing 25 RCs and 25 LHCIIs at 20% of their original size were randomly deposited on a field with a periodic boundary (**Supplementary Fig. S13)**. After resolving any overlaps, the disc size was increased by 0.08%, and a new round of fitting was started until the particle grew to 100 % of its original size. For the mixture of WT RC and LHCII proteins two parameters, the centroid coordinate $\boldsymbol{C(x,y)}$ and the orientation of the protein with respect to the reference axis ($\vartheta$), were randomly assigned and any appearing overlap was resolved with small perturbation of $C(x,y)$ and/or $\vartheta$. To recapitulate the covalent association within an RC#LHCII chimera two additional parameters were introduced that described the distance ($d$) and relative angle ($\varphi$) of the two proteins. Parameter $d$ was varied in the permitted range determined by the flexible linkers on the two proteins and the SpyCatcher/SpyTag complex (0 to 2.9 nm). The initial configuration was relaxed with small steps (0.02 of RC maximal diameter) and the self-correlation was assessed by the self-dynamic overlap function $Q(a, \tau)$ converged following **Eq. S1**[3]:

$$Q_s(a,\tau) = \left\langle \frac{1}{N}\sum_{i=1}^{N_P}\exp(-\frac{[\mathbf{r}_i(\tau)-\mathbf{r}_i(0)]^2}{a^2})\right\rangle \quad \textbf{(S1)}$$

where $\boldsymbol{r}_i$ is the coordinate(x,y) of each object at time $\tau$. Each simulation condition was repeated ten times to capture the ensemble of the system given that each setup had 50 objects. The simulations were conducted at various surface occupations ranging from 5% to 60% to understand the consequences for energy flow within mixtures of independent RC and LHCII proteins and within and between the LHCII#RC chimeras. The selected average self-correlation – $Q_s(a,\tau)$ of all proteins is shown in **Supplementary Fig. S20** with the assignment of $a$ equal to 1. Representative final states of the simulations are shown in **Supplementary Fig. S14**. The centre-centre distances of all possible pairs of RCs and LHCIIs ($R$) were extracted and used to calculate $E_{sim}$ based on a FRET mechanism, as shown in **Supplementary Fig. S13** and described below.

**Section 4 – Estimation of ET based on FRET mechanism**

Based on simulated centre-to-centre distances the overall ET efficiency between LHCII energy donors and RC energy acceptors under various conditions could be determined using the classic FRET equation:

$$R_o^6 = 8.785\times 10^{-25}\frac{\Phi_D\kappa^2 J}{n^4} \quad (in\ cm^6) \quad \textbf{(S2)}$$

which describes $R_o$, the separation distance of a single donor-acceptor pair at 50 % ET efficiency[4]. The quantum yield ($\Phi_D$) of LHCII was measured to be 0.14 (**Supplementary Fig. S15** & **Supplementary Table S1**), which was consistent with literature values[5–7]. The spectral overlap ($J$) between the LHCII fluorescence and RC absorbance spectra was estimated to be 9.36 x $10^{-13}$ $cm^{-3}M^{-1}$ as summarized in **Supplementary Table S1**. The refractive index $n$ was set to that of water (1.33) and the orientation factor $\kappa^2$ was assumed to be 2/3. To account for the considerations that a previously observed effect of nano-structured silver is to increase the emission output of LHCs by light scattering and plasmon resonance[8] and the possibility of better alignment of pigment dipoles due to adsorption to the surface, the calculated $R_o^6$ was multiplied with a correction factor equal to 3.5 to bring the result of the simulation to the level of the experimental observations.

The $E_{DA}$ of every possible pair of RC and LHCII was then calculated using:

$$E_{DA}(i) = \frac{R_o^6}{R_o^6 + R^6(i)} \quad (i = 1,2,...,N_{RC}) \quad \textbf{(S3)}$$

where $R(i)$ is the separation distance between each LHCII and every RC ($i = 1,2,\ldots,N_{RC}$) within the simulation field (**Supplementary Fig. S13**). Because efficiency of ET is the consequence of competition of different relaxation pathways for the same pool of excitons (LHCII excitation states), the $E_{DA}$ for a single donor-acceptor pair could be written as the ratio of relaxation kinetic rates according to:

$$E_{DA}(i) = \frac{k_{ET}(i)}{k_{other} + k_{FL} + k_{ET}(i)} \quad (i = 1,2,...,N_{RC}) \quad \textbf{(S4)}$$

where the rates of all non-fluorescent relaxation processes were represented by $k_{others}$, that of the fluorescent pathway was denoted as $k_{FL}$, and resonant energy transfer was indicated by $k_{ET}$ *(i)* for all possible RC ($i$ = $1,2,\ldots,N_{RC}$) from a single LHCII.

The apparent efficiency of ET could then be determined as shown in **Eq. S5**, where transfer rates between individual LHCIIs and RCs were summed to estimate the overall ET efficiency from a LHCII to all possible RCs. In this equation the apparent ET efficiency is the sum of the kinetics of all single D-A pair rates divided by the sum of all possible relaxation kinetics.

$$E_{app} = \frac{\sum_{i=1}^{N_{RC}} k_{ET}(i)}{k_{other} + k_{FL} + \sum_{i=1}^{N_{RC}} k_{ET}(i)} \qquad (i = 1, 2, ..., N_{RC}) \tag{S5}$$

Rearrangement of **Eq. S4** then gave an expression for $k_{ET}$ as a function of $E_{DA}$ :

$$k_{ET}(i) = \frac{E_{DA}(i)}{1 - E_{DA}(i)}(k_{other} + k_{FL}) \qquad (i = 1, 2, ..., N_{RC}) \tag{S6}$$

Substitution of $k_{ET}$ in **Eq. S5** with **Eq. S6** gave an expression for the apparent ET efficiency in terms of all possible single donor-acceptor pair ET efficiencies, $E_{DA}$**:**

$$E_{app} = \left\langle (\sum_{i=1}^{N_{RC}} \frac{E_{DA}(i)}{1 - E_{DA}(i)}) \frac{1}{1 + \sum_{i=1}^{N_{RC}} \frac{E_{DA}(i)}{1 - E_{DA}(i)}} \right\rangle \tag{S7}$$

**Eq. S7** provided an estimation of ET efficiency that could be used to understand the experimental observations on ET in solution and on densely-packed surfaces. The values resulting from the simulation are summarized in **Fig. 4i** and discussed in the main text. It should be noted that the simulation did not aim to recreate precisely all possible interactions of RCs and LHCs but rather was aimed at capturing the relationship between packing density, inter-protein distance and ET efficiency in a simple 2D system, and revealing the impact of tethering individual FRET donors to individual FRET acceptors in such a system.

For analysis of ET in the RC#LHCI#RC chimeras with two RC acceptors ($N_{RC}$ = 2), **Eq. S7** could be written as **Eq. S8** to assess the contribution of each RC.

$$E_{FL,P870} = (\frac{E_{a1}}{1 - E_{a1}} + \frac{E_{a4}}{1 - E_{a4}}) \frac{1}{1 + \frac{E_{a1}}{1 - E_{a1}} + \frac{E_{a4}}{1 - E_{a4}}} \tag{S8}$$

Rearrangement of the right part of **Eq. S8** produced **Eq. 5** shown in the main text for distinguishing the contributions to the ET efficiency of the RC attached to Lhca1($E_{a1}$) and that attached to Lhca4 ($E_{a4}$).

### Supplementary figures

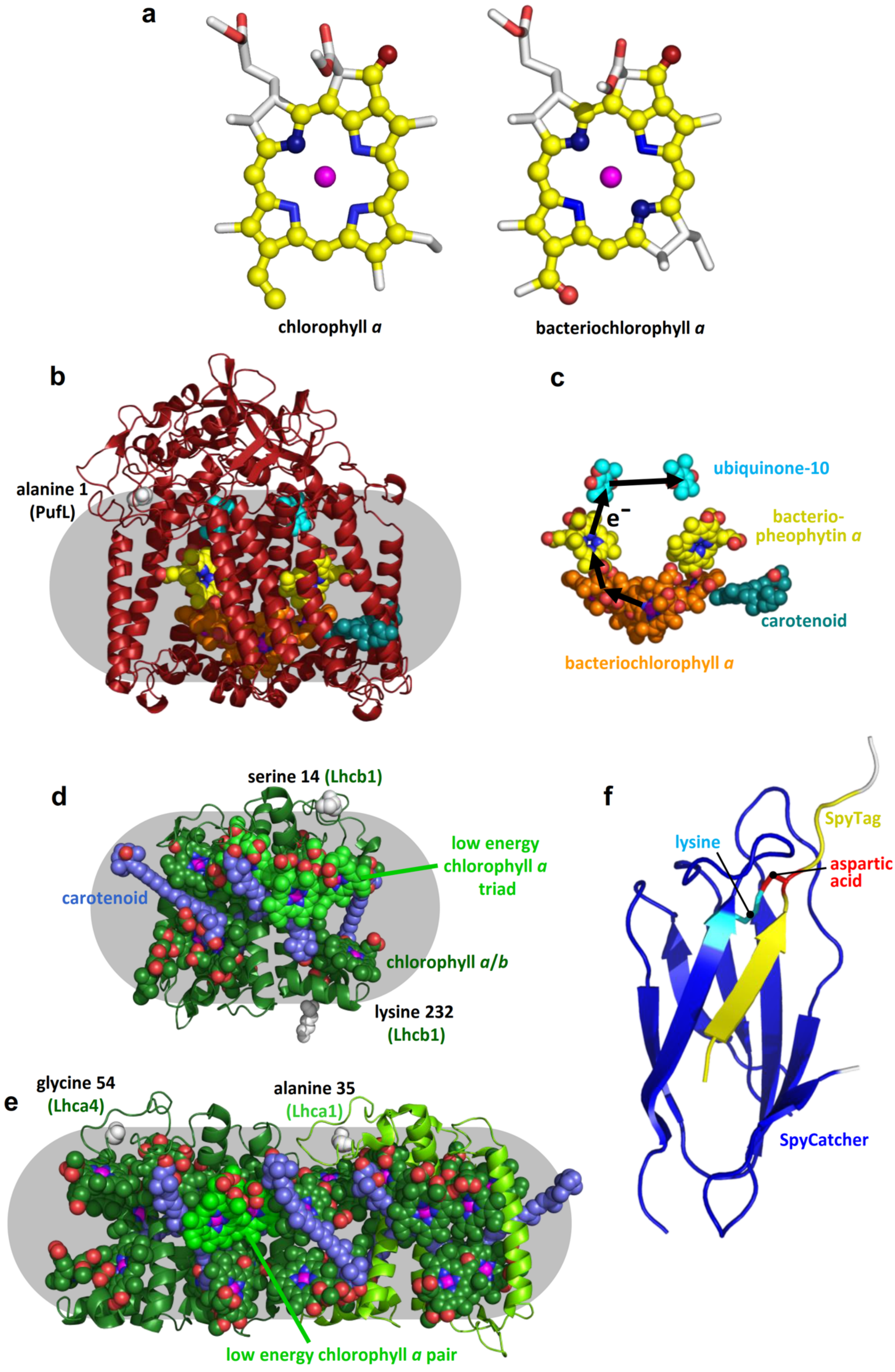

**Supplementary Fig. S1.** Structures of components and RC mechanism. **a** Structures of the macrocycles of chlorophyll *a* and bacteriochlorophyll *a*. Atoms involved in the conjugated electron system are highlighted with spheres and coloured yellow (carbon), dark red (oxygen) or dark blue (nitrogen). Other atoms are coloured white (carbon), blue (nitrogen), red (oxygen), magenta (magnesium). The extent and shape of each delocalised electron system determines the absorbance properties of the cofactor. **b** The *Rba. sphaeroides* RC comprises three polypeptides (maroon ribbons) that scaffold its cofactors. Grey shading approximates the dimensions of a detergent micelle. Non-native polypeptides were added before alanine 1 of the PufL subunit (white spheres, top left). **c** View of the structure in **b** with the protein and cofactor side chains removed. Cofactors comprise four bacteriochlorophyll *a* (orange carbons), two bacteriopheophytin *a* (yellow carbons), two ubiquinone-10 (cyan carbons) and one carotenoid (teal carbons). Four-step charge separation (black arrows) takes place between two bacteriochlorophylls that form the P870 primary electron donor and a dissociable ubiquinone (and see Fig. 1d). **d** LHCII comprises a single Lhcb1 polypeptide (green ribbon) that scaffolds eight chlorophyll *a* and six chlorophyll *b* (green carbons) and four carotenoids (slate carbons). The carbons of three interacting chlorophyll *a* that from a low energy cluster are highlighted in light green. Non-native polypeptides were added before serine 14 or after lysine 232 (white spheres). **e** LHCI complexes formed from a heterodimer of Lhca1 and Lhca4 (light/dark green ribbons) that scaffold 36 cofactors (23 chlorophyll *a*, six chlorophyll *b* and seven carotenoids). The carbons of two interacting chlorophyll *a* that from a low energy cluster are highlighted in light green. Non-native polypeptides were added before alanine 35 of Lhca1 or glycine 54 of Lhca4 (white spheres) which are the N-terminal residues in the resolved X-ray structure. **f** SpyTag (yellow) binds to SpyCatcher (blue) and an isopeptide bond is formed between lysine (cyan) and aspartic acid (red) side chains. Residues removed from the C-terminus of SpyTag are coloured white (top right).

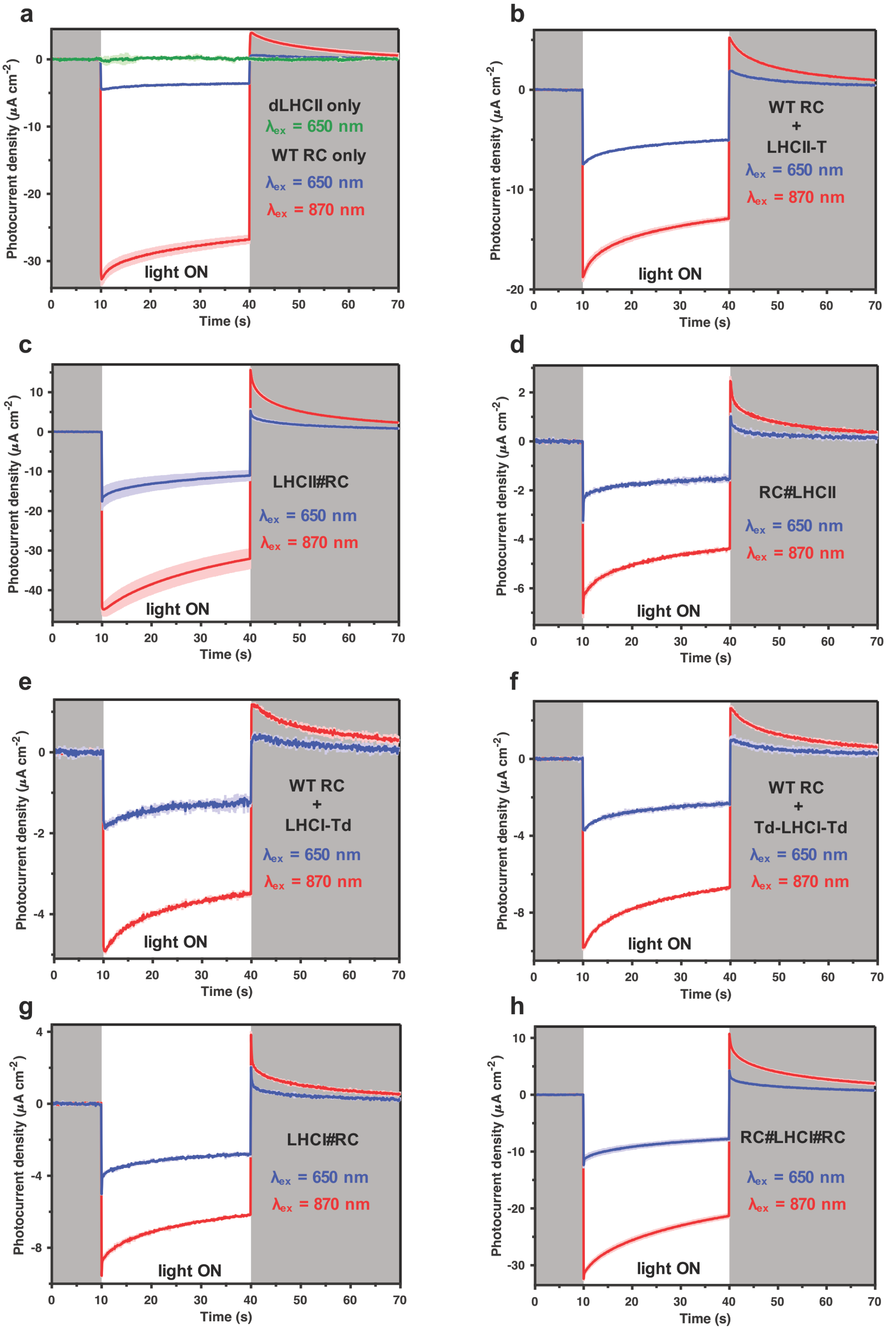

**Supplementary Fig. S2.** Photocurrents from electrodes coated with individual proteins, protein mixtures or chimeras. LED illumination was at 650 nm (power 6.7 mW cm$^{-2}$) where RCs absorb weakly but the LHCs absorb strongly, or at 870 nm (power 32 mW cm$^{-2}$) where only RCs absorb (50 nm FWHM for both LEDs). Traces are the average of three measurements with colour shading showing the standard deviation. Panel **a** shows data from two electrodes coated with either LHCII or RCs.

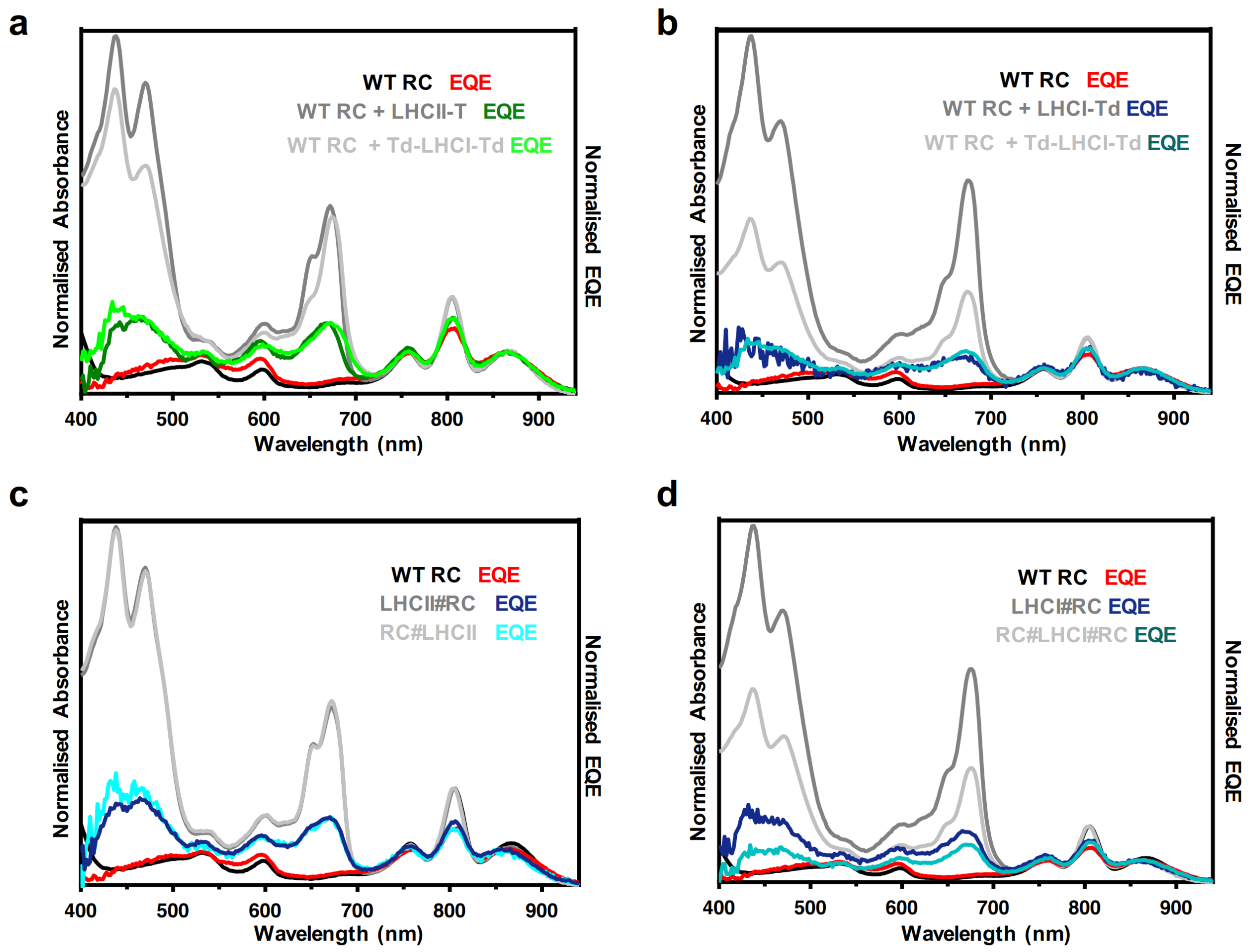

**Supplementary Fig. S3.** Normalised absorbance and EQE action spectra. EQE spectra were recorded using a tungsten light source. For comparison, absorbance spectra were normalised at the maximum of the RC band at 804 nm. Each EQE spectrum was then normalised to the associated absorbance spectrum at the maximum of the RC P870 absorbance band.

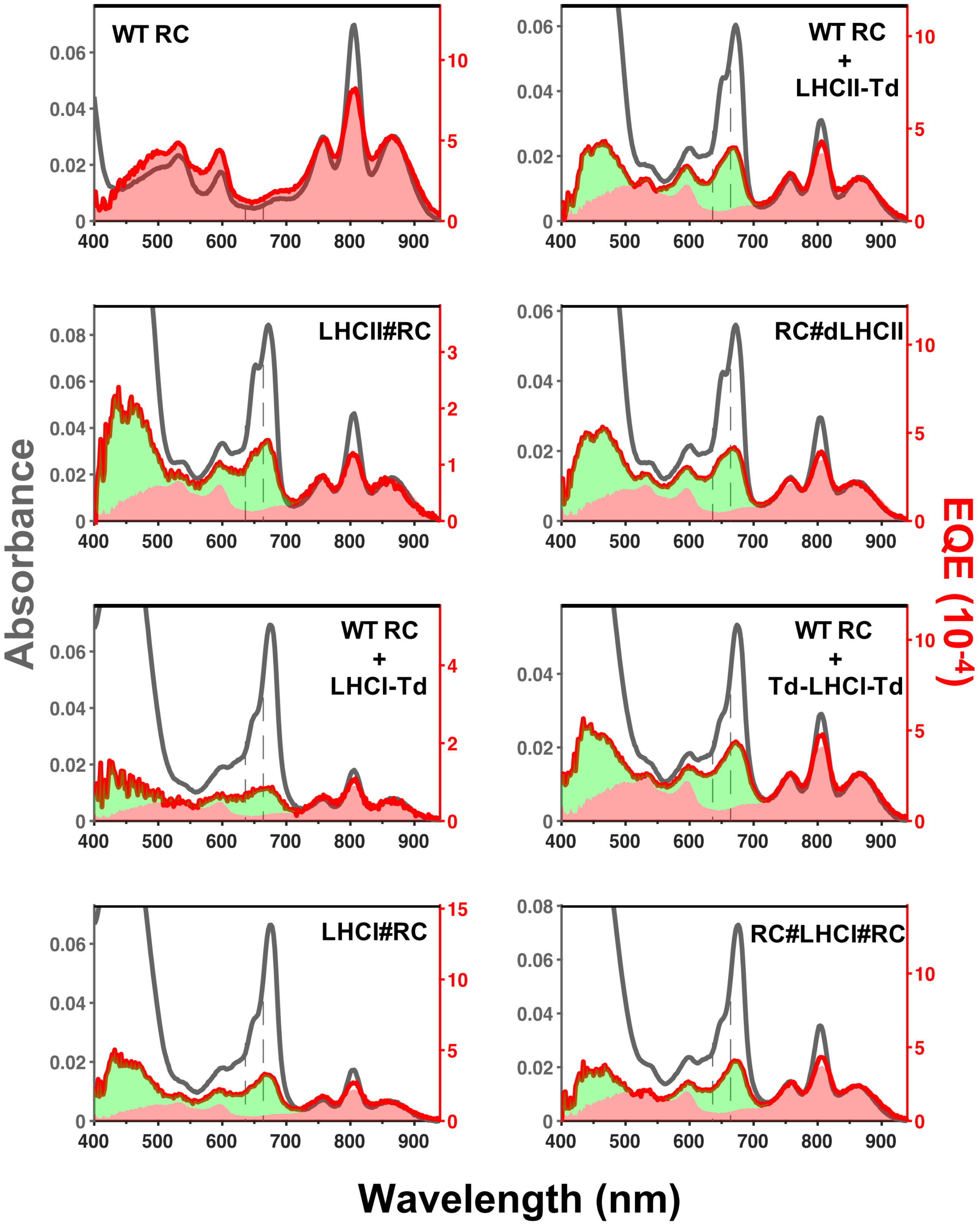

**Supplementary Fig. S4.** Absorbance (grey) and EQE action spectra (red) recorded using a tungsten light source for individual bio-photoelectrodes. Spectra were normalised at the maximum of the RC P870 absorbance band. The red and green shaded areas indicate the contributions of the RC and LHC, respectively, to the overall EQE spectrum.

**a**

```
     dLHCII                       MSPWY-Lhcb1-VPGKLEHHHHHHHHHH*
   Td-dLHCII            MGSHHHHHHSGAHIVMVDAYKGSPWY-Lhcb1-VPGKLE*
     LHCII-T                  MGSRKTVAKPKGPSGSPWY-Lhcb1-VPGKLEGAHIVMVDAYKPTKSSGHHHHHH*

    T-dLHCII         MGSHHHHHHSGAHIVMVDAYKPTKGSPWY-Lhcb1-VPGKLE*

          L1                  MGSEQKLISEEDLGTAGNV-Lhca1-IPFN*
       Td-L1        MGSEQKLISEEDLGTAHIVMVDAYKGAGNV-Lhca1-IPFN*
       Td-L4            MGSHHHHHHSGAHIVMVDAYKGKKGE-Lhca4-QTFN*
```

**b**

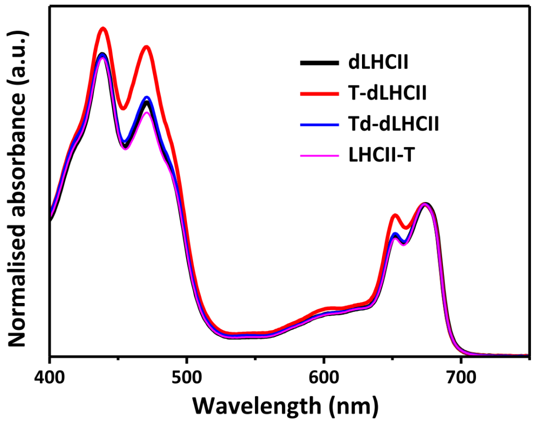

**c**

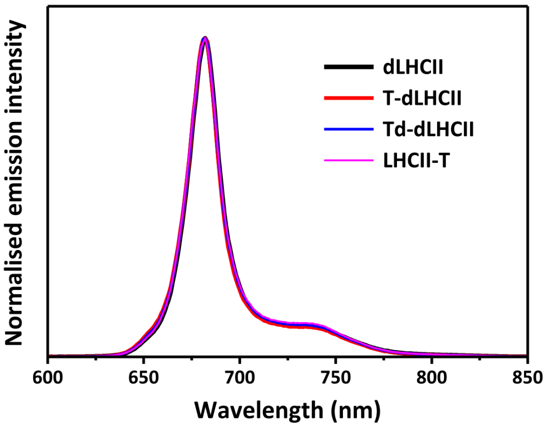

**d**

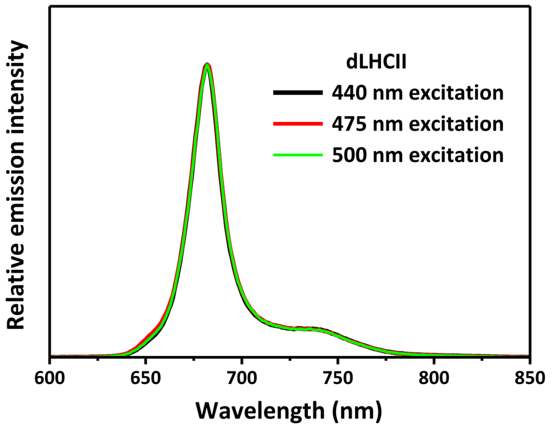

**e**

| Sample | Chl *a*/Chl *b* | Chl/[LHCII] |
|---|---|---|
| dLHCII | 1.43 | 12.51 |
| T-dLHCII | 1.09 | 13.48 |
| Td-dLHCII | 1.37 | 12.85 |
| LHCII-T | 1.50 | 11.96 |

**Supplementary Fig. S5.** LHC modification and LHCII refolding. **a** LHC protein sequences are marked in bold, with the dispensable sequence at the N-terminus of Lhcb1 highlighted in grey. Added His and Myc (EQKLISEEDL) tags are highlighted in cyan, the full-length and truncated version of SpyTag are highlighted in yellow, and linkers are underlined. Only the first four and last four residues of each LHC protein are shown. Data for construct T-dLHCII were essentially identical to those for Td-dLHCII and are not discussed in the main text. **b** Absorbance spectra of the four refolded LHCII variants, normalised at the maximum of the low energy chlorophyll *a* band. **c** Normalised emission spectra of these complexes obtained with 440 nm excitation. **d** Normalised emission spectra for dLHCII illustrating that emission profile was independent of excitation wavelength, indicating correct refolding. **e** Deduced cofactor compositions for each refolded LHCII.

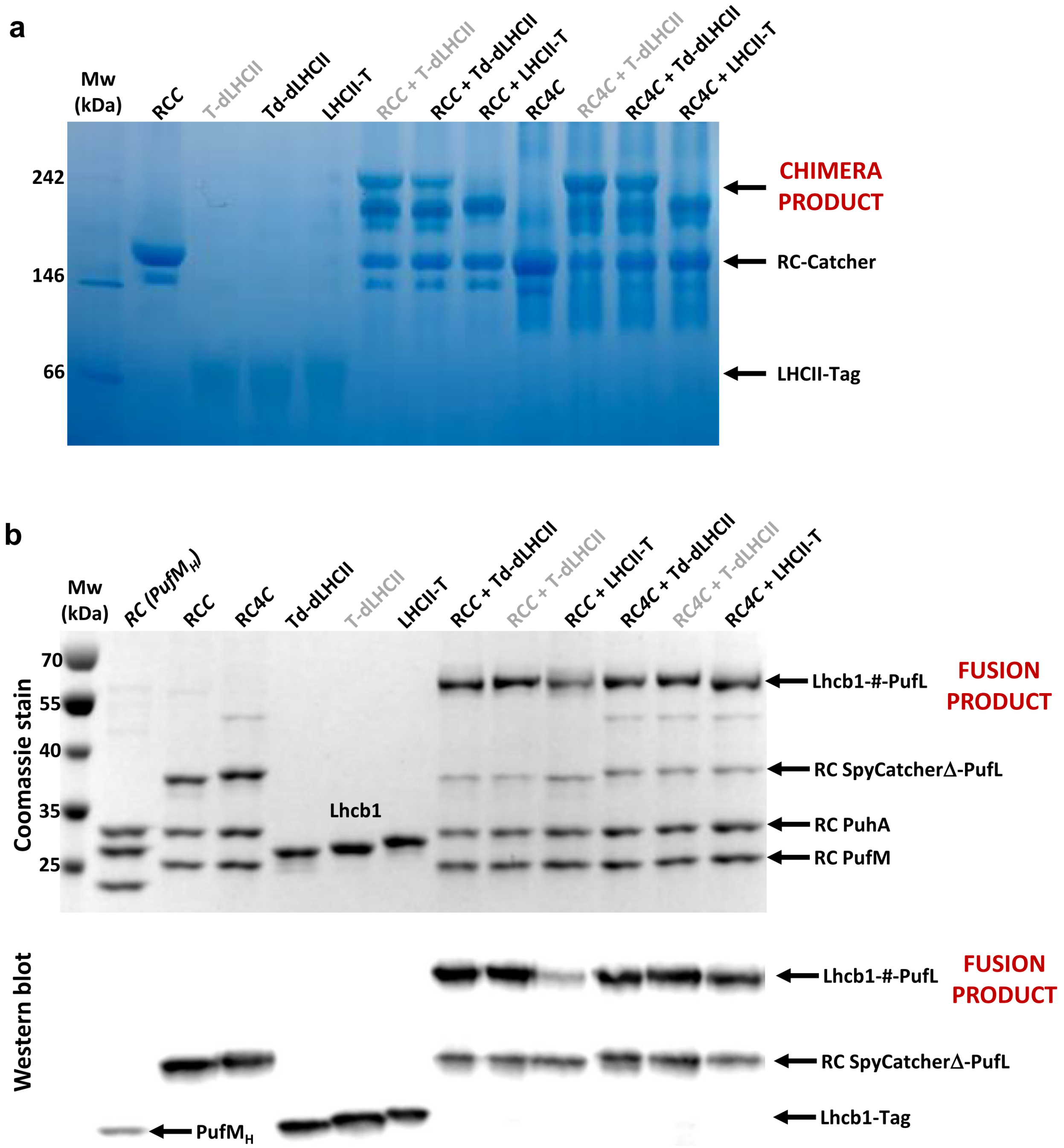

**Supplementary Fig. S6.** Formation of covalent RC-LHCII chimeras and fusion proteins. Gels and blots show the outcome of mixing a two-fold excess of SpyCatcherΔ-adapted RCs with SpyTag(Δ)-adapted LHCII proteins. Lanes labelled in grey show proteins/combinations involving variant T-dLHCII that are not referred to in the main text. **a** Blue native PAGE of individual proteins and mixtures. Mixing of SpyCatcherΔ-adapted RCs (bands labelled RC-Catcher) with SpyTag or SpyTagΔ-adapted LHCII complexes (bands labelled LHCII-Tag) produces higher molecular weight chimera products. **b** SDS PAGE and anti-His western blotting of individual proteins and mixtures. The RC is made up from three polypeptides, PufL (L-polypeptide), PufM (M-polypeptide), PuhA (H-polypeptide). Mixing of SpyCatcherΔ-adapted RCs with SpyTag- or SpyTagΔ-adapted LHCII complexes produces a Lhcb1-#-PufL fusion polypeptide (labelled FUSION PRODUCT) where # denoted the SpyCatcher/Tag sequences covalently locked by an isopeptide bond. Western blotting detected the His-tag on the Lhcb1-#-PufL fusion polypeptide and the His-tags on unreacted RC SpyCatcherΔ-PufL and residual Lhcb1-Tag variants.

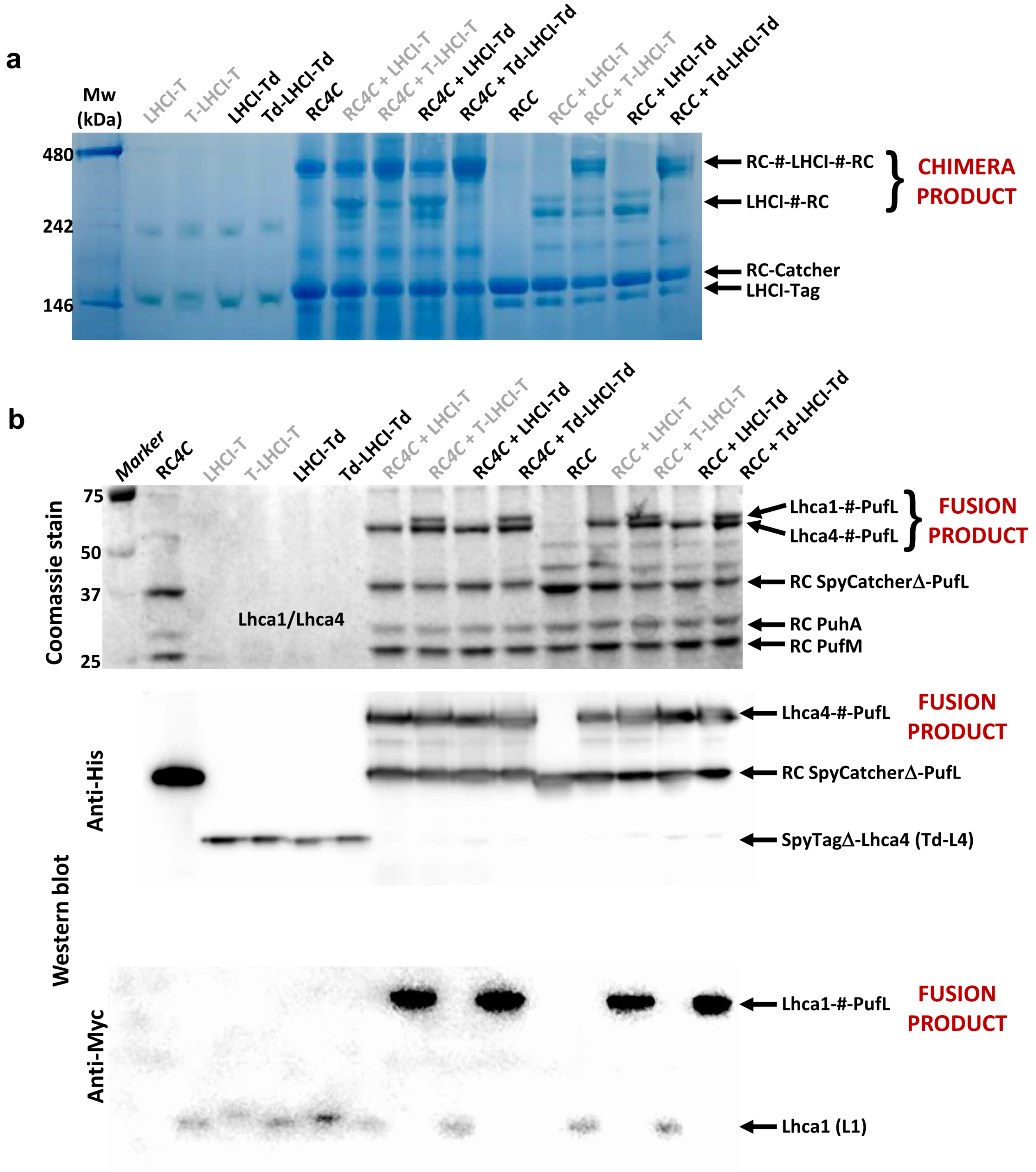

**Supplementary Fig. S7.** Formation of covalent RC-LHCI chimeras and fusion proteins. Gels and blots show the effect of mixing a three-fold excess of SpyCatcherΔ-adapted RCs with single or doubly SpyTag(Δ)-adapted LHCI complexes. In addition to combinations described in the text, data are shown for an equivalent set of constructs in which LHCI was either singly (LHCI-T) or doubly (T-LHCI-T) modified with the full length SpyTag peptide (lane labels shown in grey), which produced identical results. **a** Blue native PAGE of individual proteins and mixtures. Mixing of SpyCatcherΔ-adapted RCs (labelled RC-Catcher) with singly or doubly SpyTag- or SpyTagΔ-adapted LHCI complexes (labelled LHCI-Tag) produces higher molecular weight products denoted as LHCI-#-RC or RC-#-LHCI-#-RC (labelled CHIMERA PRODUCT) where # is the SpyCatcher/Tag domain. The multiple bands seen for these high molecular weight products may have arisen from conformational heterogeneity. **b** SDS PAGE and western blotting of individual proteins and mixtures revealing the formation of covalently locked Lhca4-#-PufL and/or Lhca1-#-PufL fusion proteins. LHCI is a heterodimer of Lhca1 and Lhca4. Mixing of SpyCatcherΔ-adapted RCs with singly SpyTag- or SpyTagΔ-adapted LHCI complexes (on Lhca4) produces a covalently linked Lhca4-#-PufL polypeptide (labelled FUSION PRODUCT) where # is the SpyCatcher/Tag sequences connected by an isopeptide bond. Mixing of SpyCatcherΔ-adapted RCs with doubly SpyTag- or SpyTagΔ-modified LHCI complexes produces covalently linked Lhca1-#-PufL and Lhca4-#-PufL polypeptides (labelled FUSION PRODUCTS). Western blotting with anti-His antibodies detected the His-tag on the Lhca4-#-PufL fusion peptide and the His-tags on unreacted RC SpyCatcherΔ-PufL and residual Lhca4 Td-L4 polypeptide. Western blotting with anti-Myc antibodies detected the Myc-tag on the Lhca1-#-PufL fusion peptide and the Lhca1 L1 polypeptide.

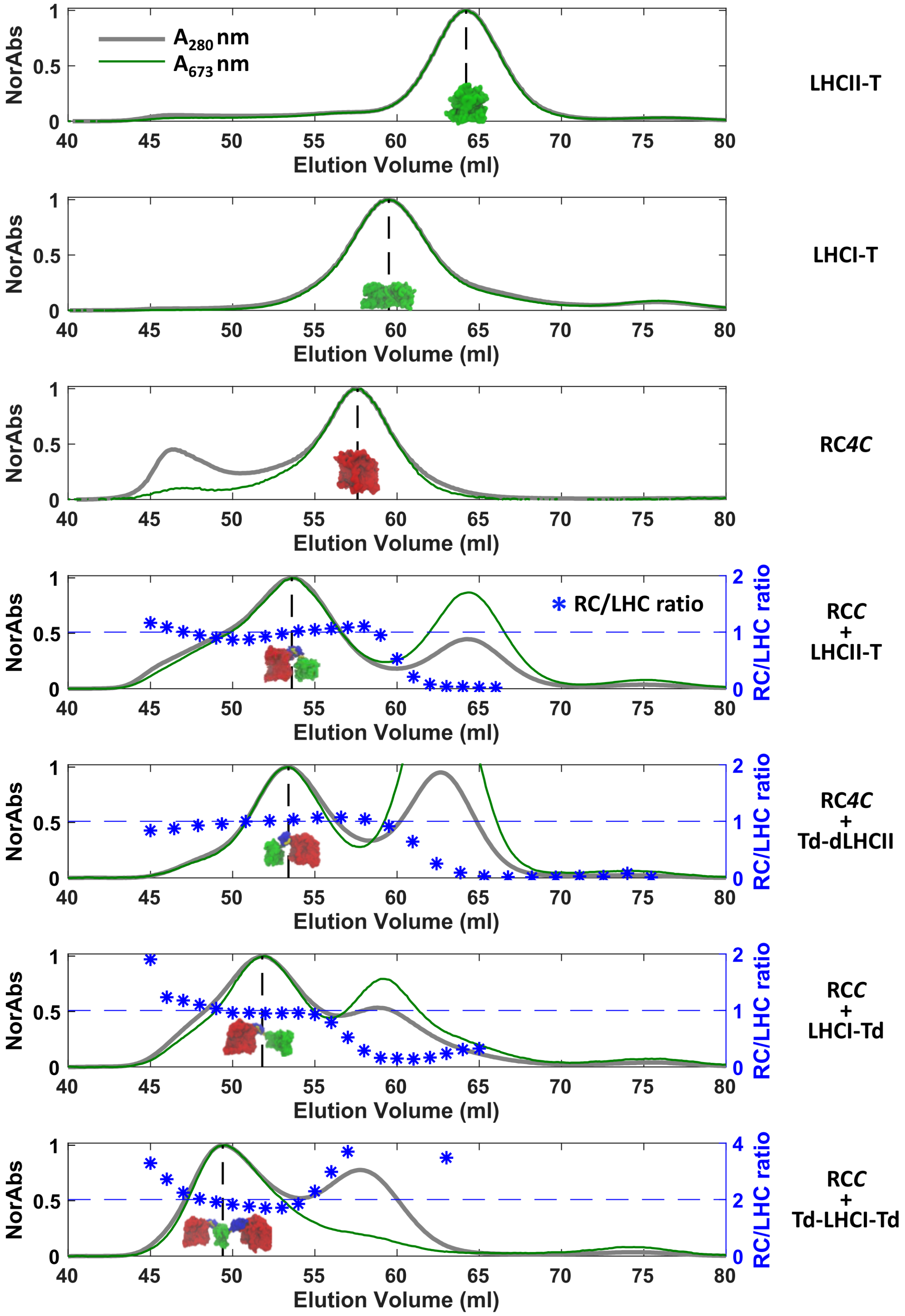

**Supplementary Fig. S8.** Purification of individual proteins or chimeras by size-exclusion chromatography. Column elution traces monitoring total protein absorbance at 280 nm (grey) and pigment-protein absorbance at 672 nm (green). Traces were normalized at the product band of interest. For all chimeras the RC/LHCII molar ratio of each collected fraction was determined from its absorbance spectrum.

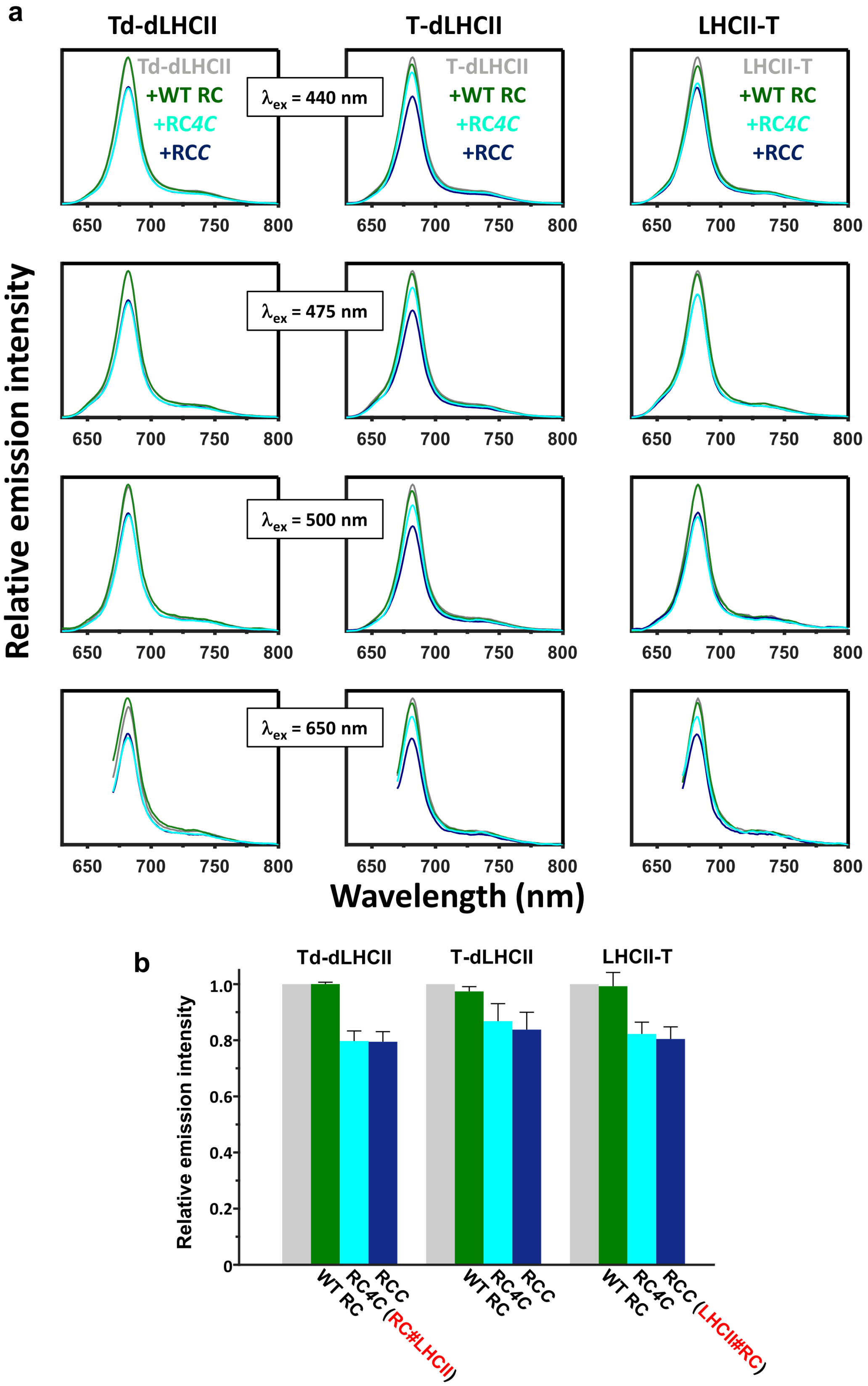

**Supplementary Fig. S9.** Emission quenching in RC-LHCII chimeras. **a** Spectra recorded with four excitation wavelengths for a control and three mixtures with a 2:1 RC:LHCII composition. Emission by each LHCII (grey spectra) was not significantly quenched by adding WT RCs (green spectra), but was reduced on adding either of the SpyCatcherΔ-adapted RCs (cyan/blue spectra) due to formation of chimeras. **b** LHCII emission at 681 nm from a control and three mixtures with a 2:1 RC:LHCII composition. Emission by each LHCII (grey) was not significantly quenched by adding WT RCs (green), but was reduced on adding either of the SpyCatcherΔ-adapted RCs (cyan/blue) due to formation of chimeras. The two chimeras studied in depth are labelled in red. Error bars indicate standard deviations from two biological repeats with three technical repeats within each biological independent sample.

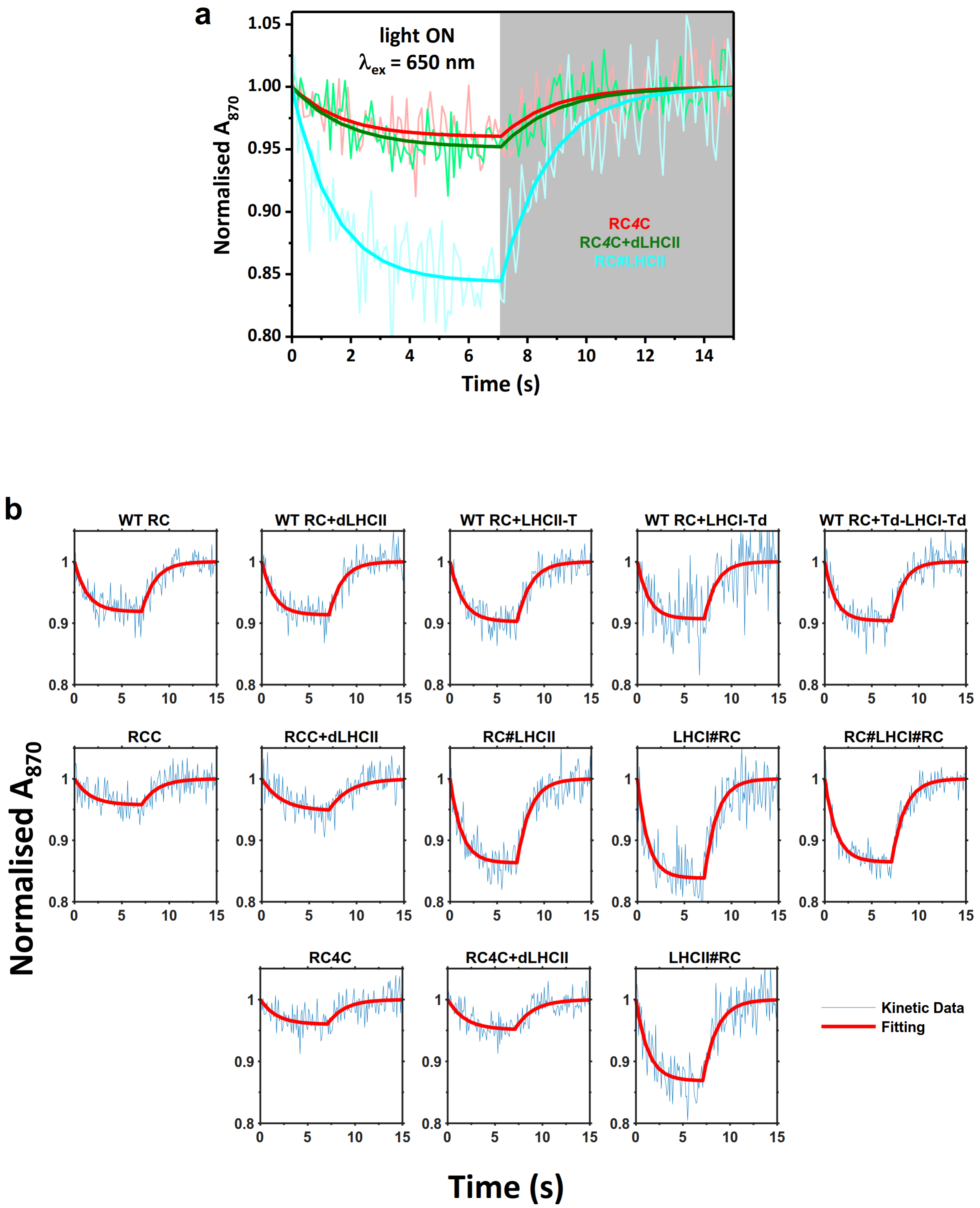

**Supplementary Fig. S10.** Photo-oxidation of P870 in RCs in response to 650 nm excitation. **a** Data and fits for photobleaching and dark recovery of P870 absorbance in RC*4C*, a 1:1 RC*4C* + dLHCII mixture and the RC#LHCII heterodimer formed by mixing RC*4C* and Td-dLHCII. **b** Data (blue) and fits (red) for all P870 bleaching experiments. Kinetic constants from the fits, carried out as described in Materials and Methods, are summarised in Supplementary Table S3.

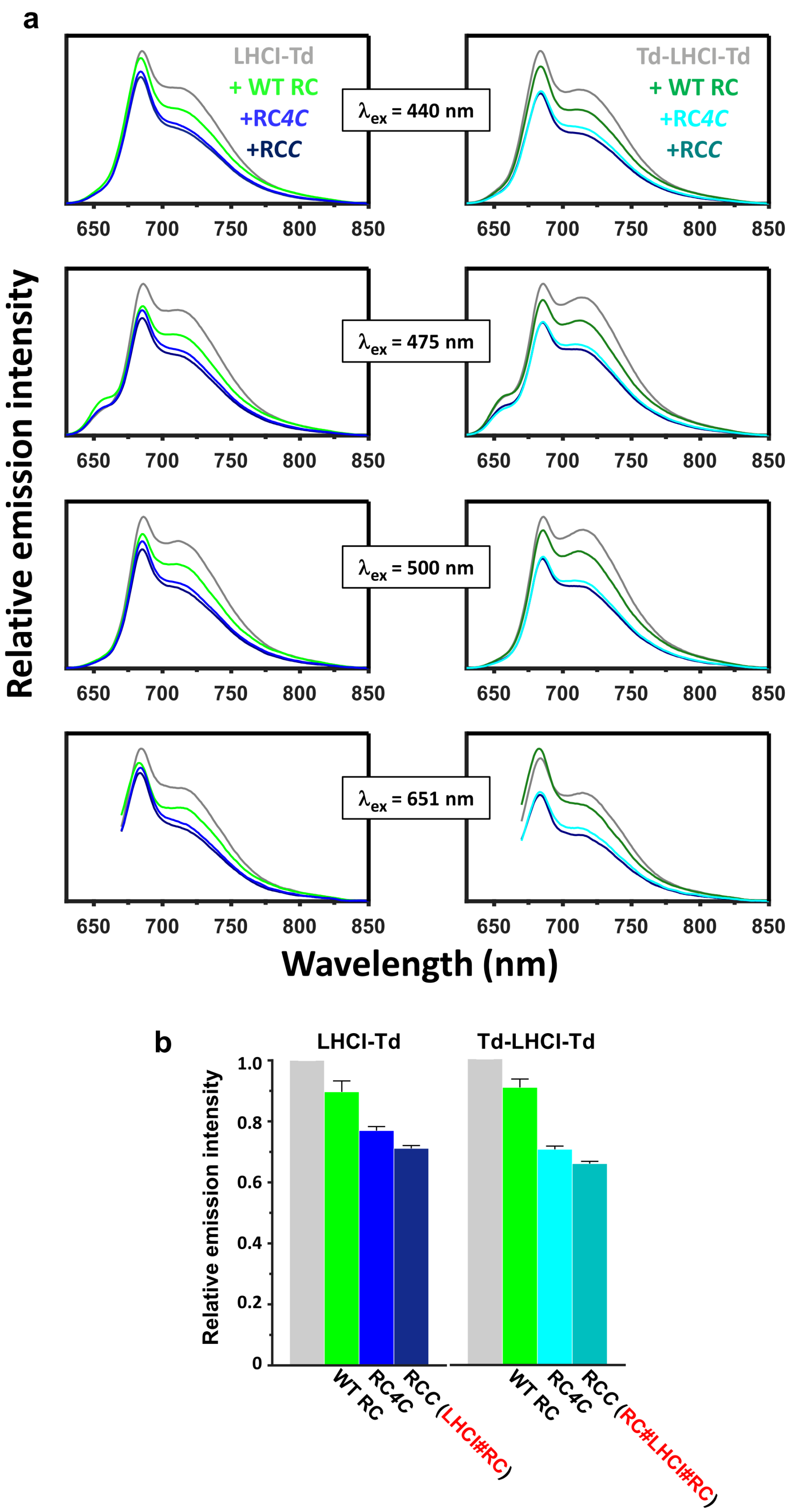

**Supplementary Fig. S11.** Emission quenching in RC-LHCI chimeras. **a** Spectra recorded with four excitation wavelengths for a control and three mixtures with a 3:1 RC:LHCI composition. Emission by each LHCI (grey spectra) was somewhat reduced on adding WT RCs (green spectra), and reduced further on adding either of the SpyCatcherΔ-adapted RCs (spectra in shades of blue) due to formation of chimeras. **b** Total LHCI emission integration from a control and three mixtures with a 3:1 RC:LHCI composition. Emission by each LHCI (grey) was somewhat reduced on adding WT RCs (green), and reduced further on adding either of the SpyCatcherΔ-adapted RCs (shades of blue) due to formation of chimeras. The two chimeras studied in depth are labelled in red. Error bars indicate standard deviations from two biological repeats with three technical repeats within each biological independent sample.

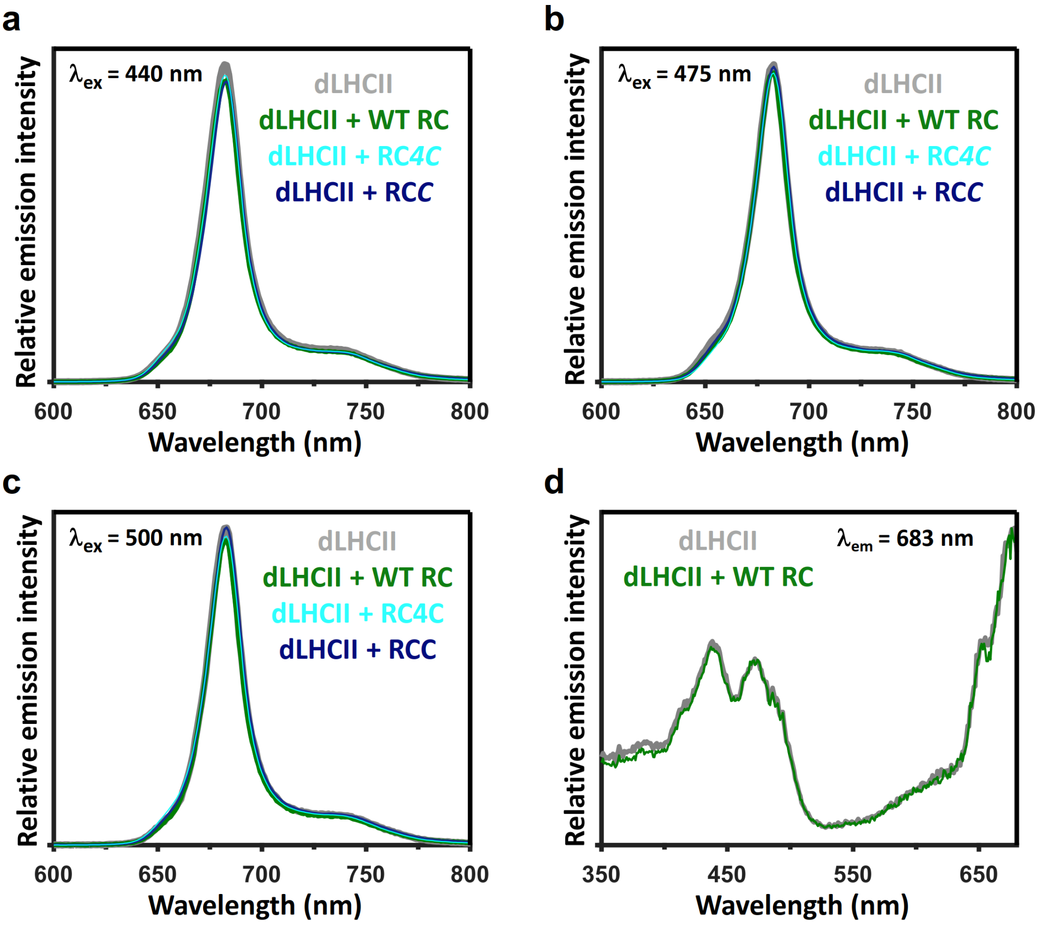

**Supplementary Fig. S12.** Emission from dLHCII is not quenched on mixing with RCs. **a-c** Emission from dLHCII with or without the WT RC, RC*4*C or RCC after correction for dLHCII absorption at the indicated excitation wavelength. **d** Excitation spectra of dLHCII emission at 683 nm. No significant differences in line shape could be observed in the presence of WT RCs.

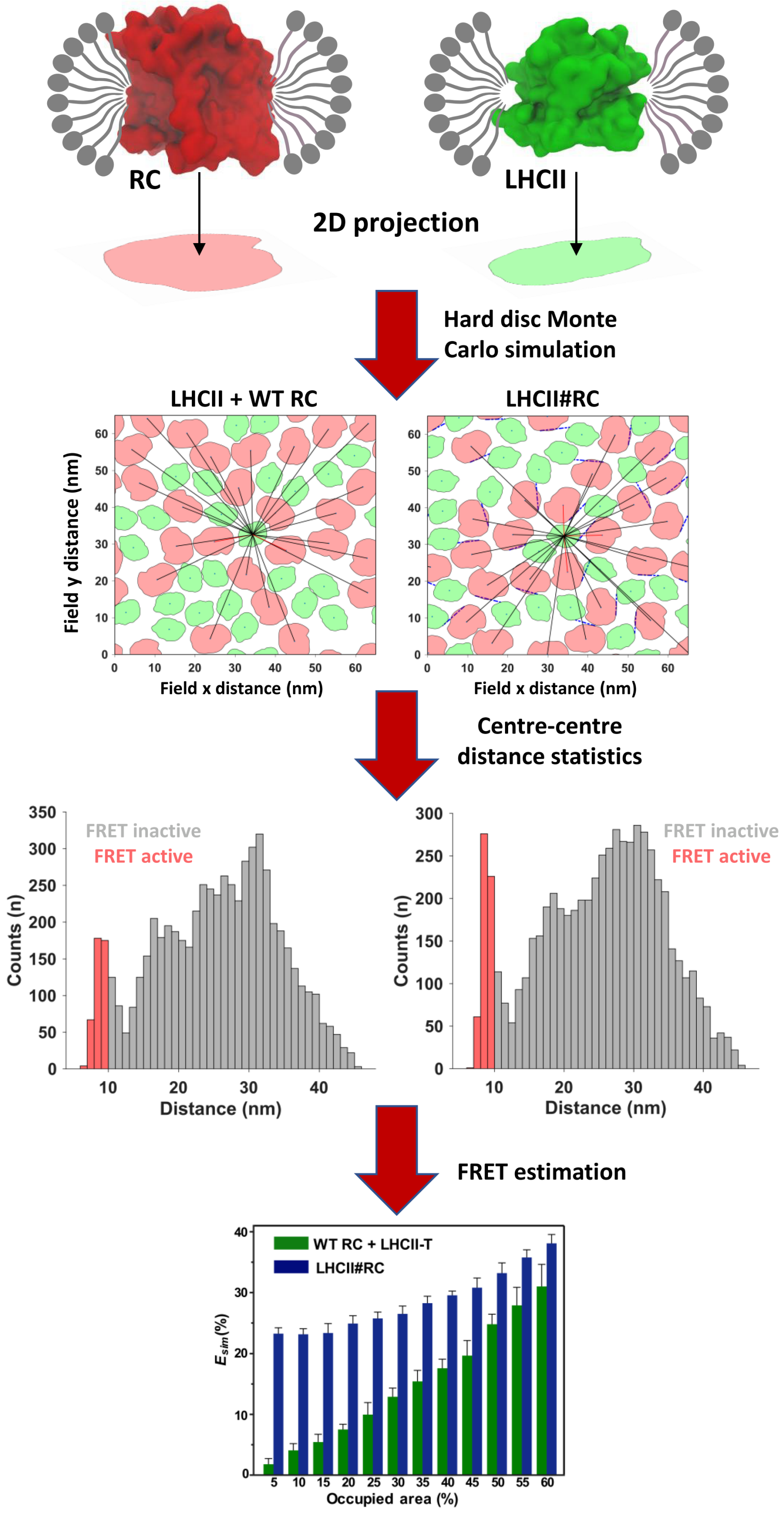

**Supplementary Fig. S13.** Workflow for 2D hard disc Monte Carlo simulation. The packing simulation was carried out with a projection of each protein/DDM complex. Centre-to-centre distances between each LHCII (green disc) and all RCs (red discs) in each simulation end configuration were determined. Donor-acceptor pairs within 10 nm in distance histogram plots (shaded red) were considered for the FRET analysis. This simulation produced a plot of apparent ET efficiency ($E_{sim}$) as a function of the percentage of the simulation area occupied by protein (also shown in Fig. 4i). Error bars were the standard deviation of ten evaluated simulations.

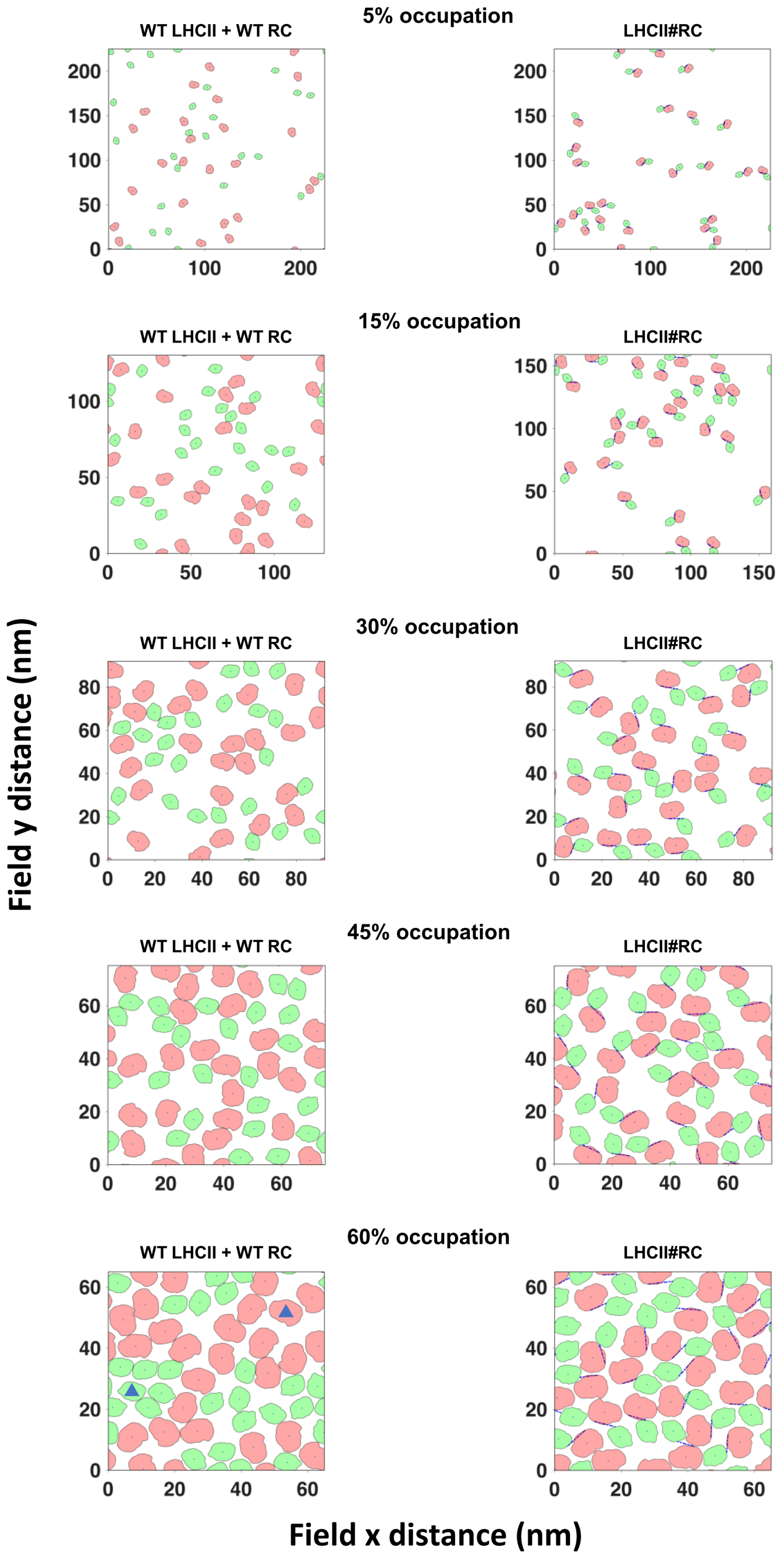

**Supplementary Fig. S14.** Representative examples of the configurations of protein complexes at equilibrium at five percentage occupancies. The centroid of each protein disc is indicated by a dot and the linkage between covalently locked LHCII#RC chimeras is signified by a dash-dot line. Occasionally at high surface occupation (60%) RCs and LHCIIs segregated into separate sub-domains such that proteins in the middle of a sub-domain were outside the FRET distance (marked with blue triangles).

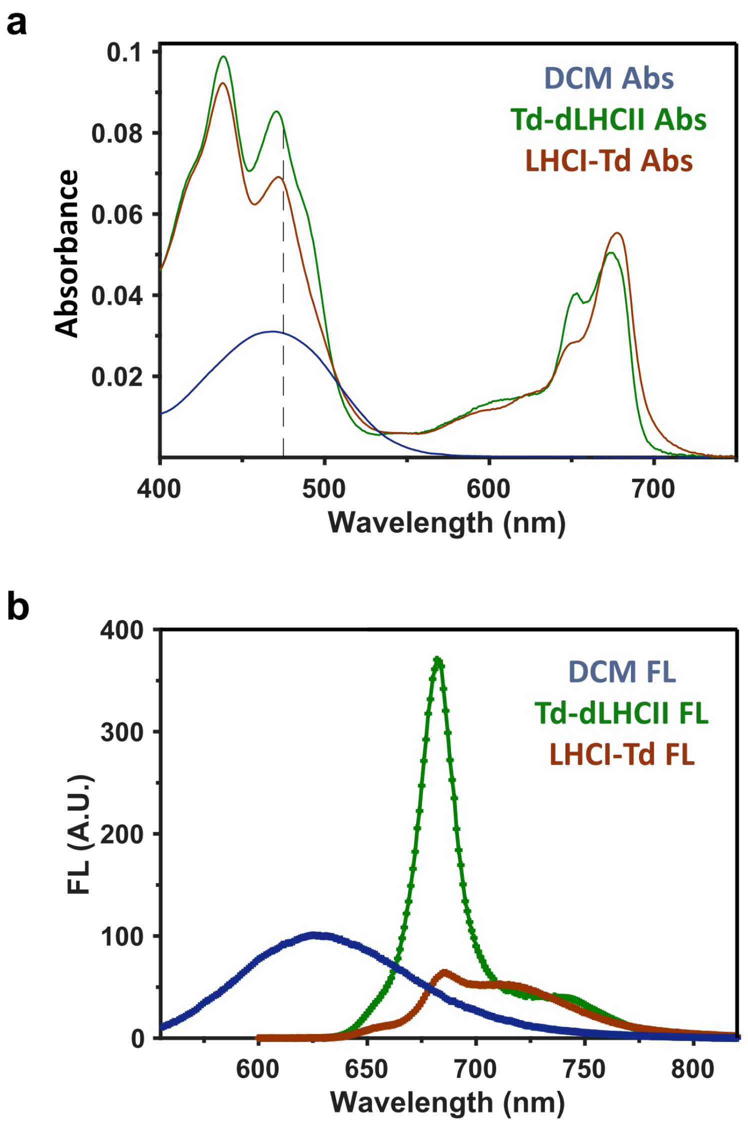

**Supplementary Fig. S15.** Relative quantum yield determination. **a** absorbance of the reference dye DCM in ethanol and the Td-dLHCII and LHCI-Td proteins in Tris/DDM buffer. The dashed line indicates the excitation wavelength at 475 nm. **b** Emission spectra of the samples shown in **a** after emission spectral response calibration.

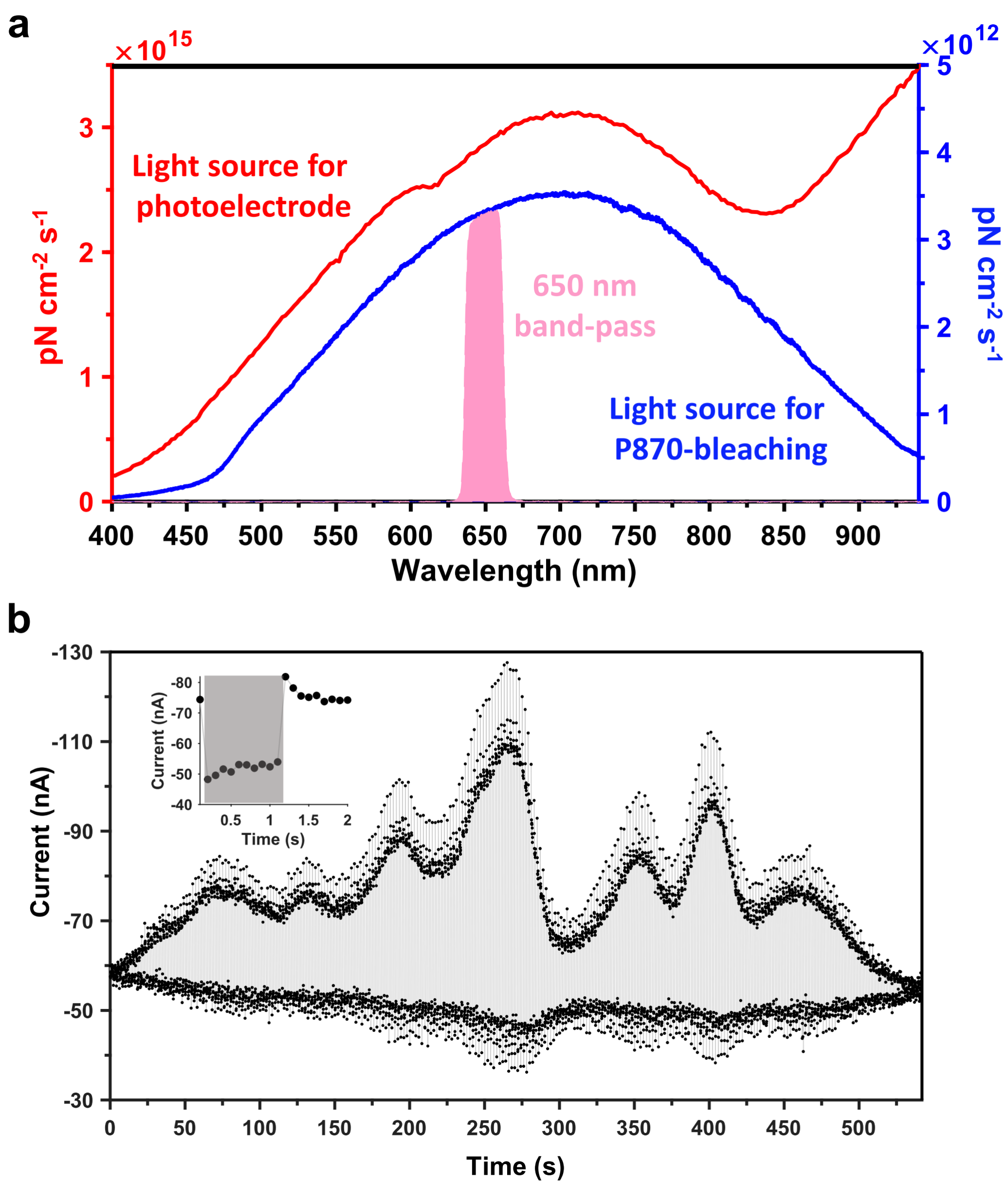

**Supplementary Fig. S16.** Recording of EQE action spectra for bio-photoelectrodes. **a** Output spectra for the tungsten lamp used to record EQE action spectra (red) and the HL-2000 lamp used for photobleaching of P870 without (blue) and with (pink) passage through a 650 nm band-pass filter. **b** Example of a raw EQE action spectrum recorded continuously over ~550 s. Each light-on/light-off cycle was as indicated in the inset with the grey shadow highlighting the period of illumination. The excitation wavelength was stepped during each dark phase. Averaging of the stable reading in each light and dark phase at each wavelength was used to compute the EQE action spectrum.

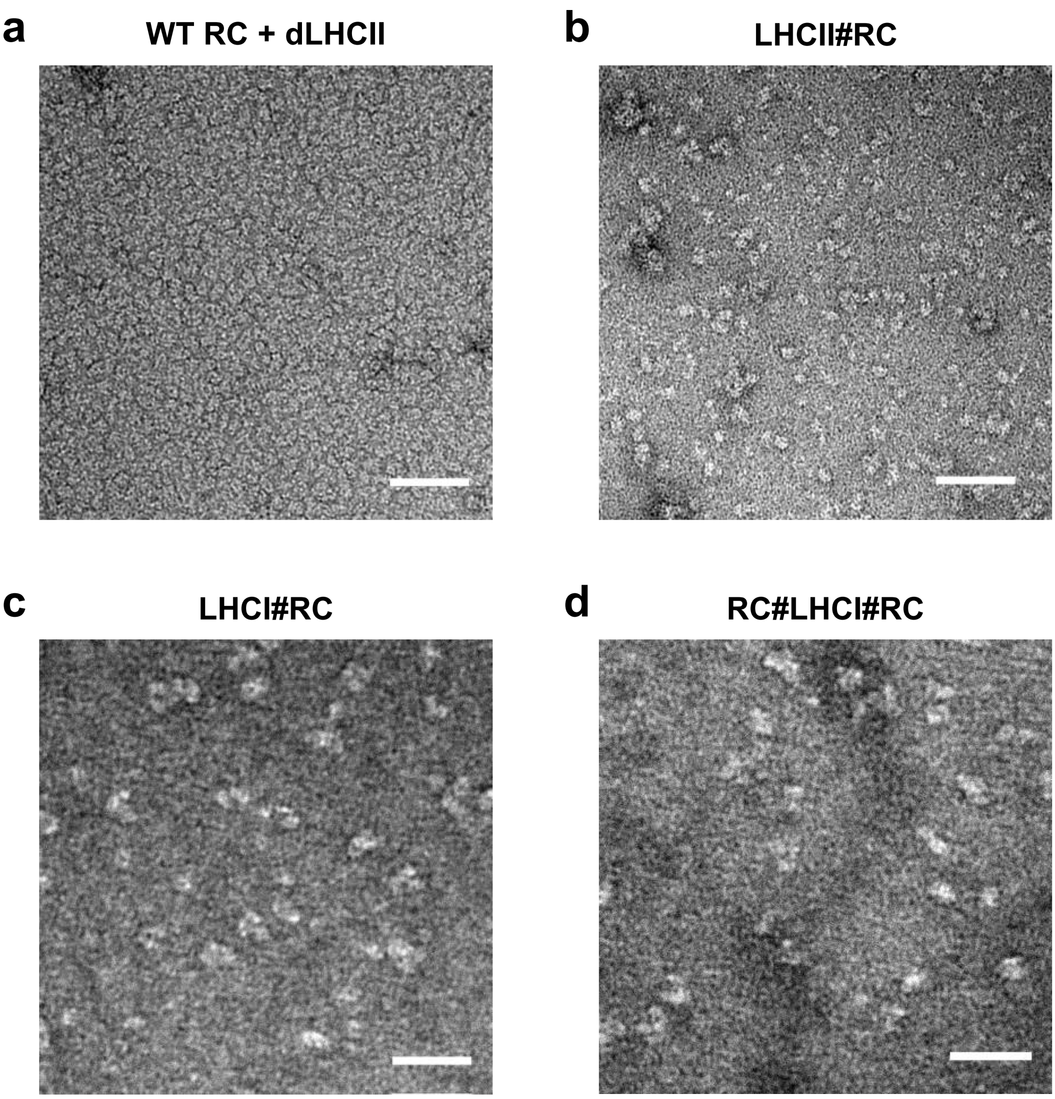

**Supplementary Fig. S17.** Negative stain TEM images. **a** Mixture of WT RC + dLHCII. **b** Heterodimeric chimera LHCII#RC. **c** Heterotrimeric chimera LHCI#RC. **d** Heterotetrameric chimera RC#LHCI#RC. White scale bars indicate 50 nm.

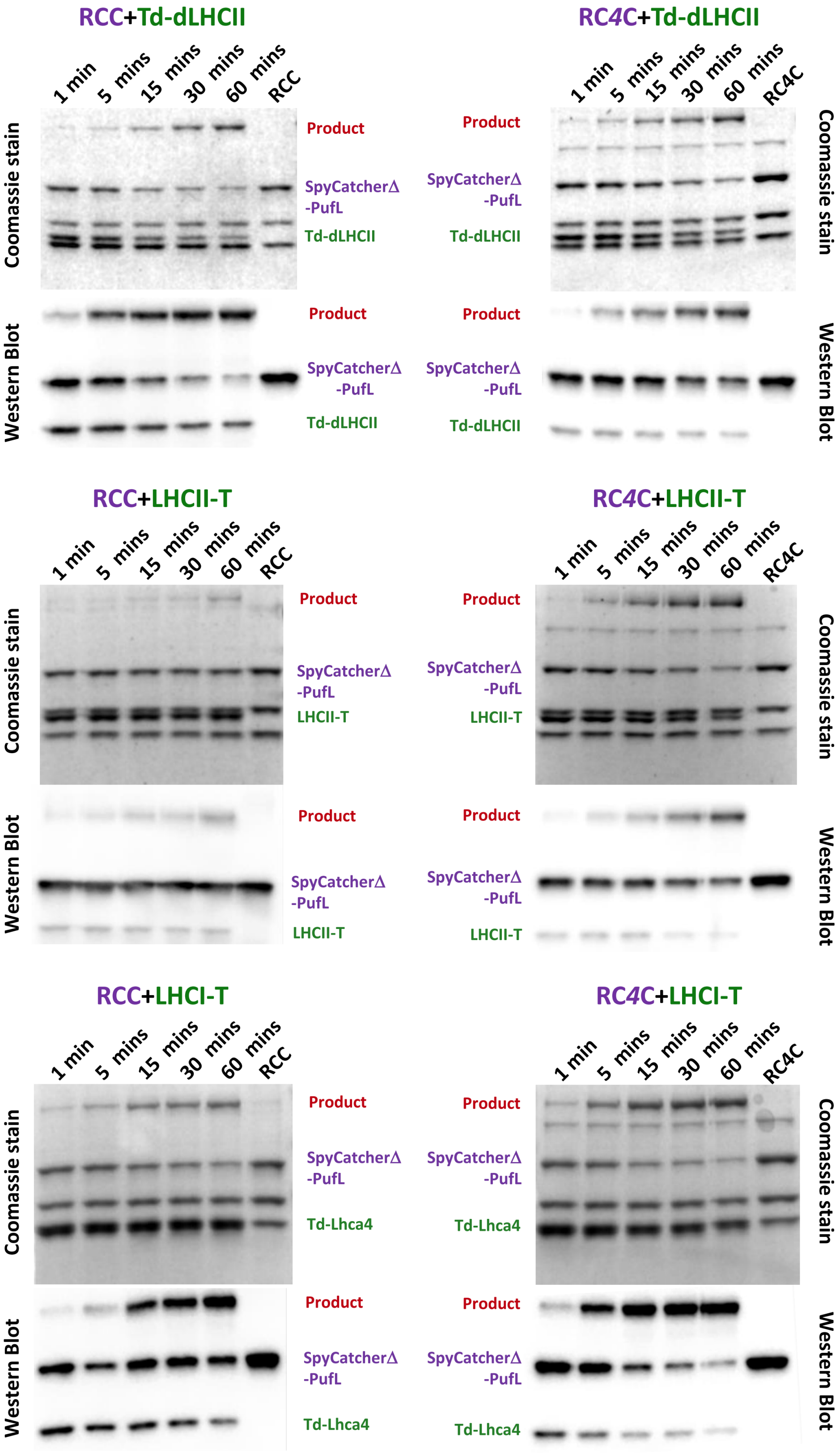

**Supplementary Fig. S18.** Characterisation of SpyCatcher/Tag reaction kinetics by SDS-PAGE and western blotting. Protein concentrations were 5 µM. The appearance of the product bands attributable to the isopeptide bond-locked fusion polypeptide was always concomitant with depletion of the reactant SpyCatcherΔ-PufL and SpyTag-LHC polypeptides, indicating covalent ligation between the two.

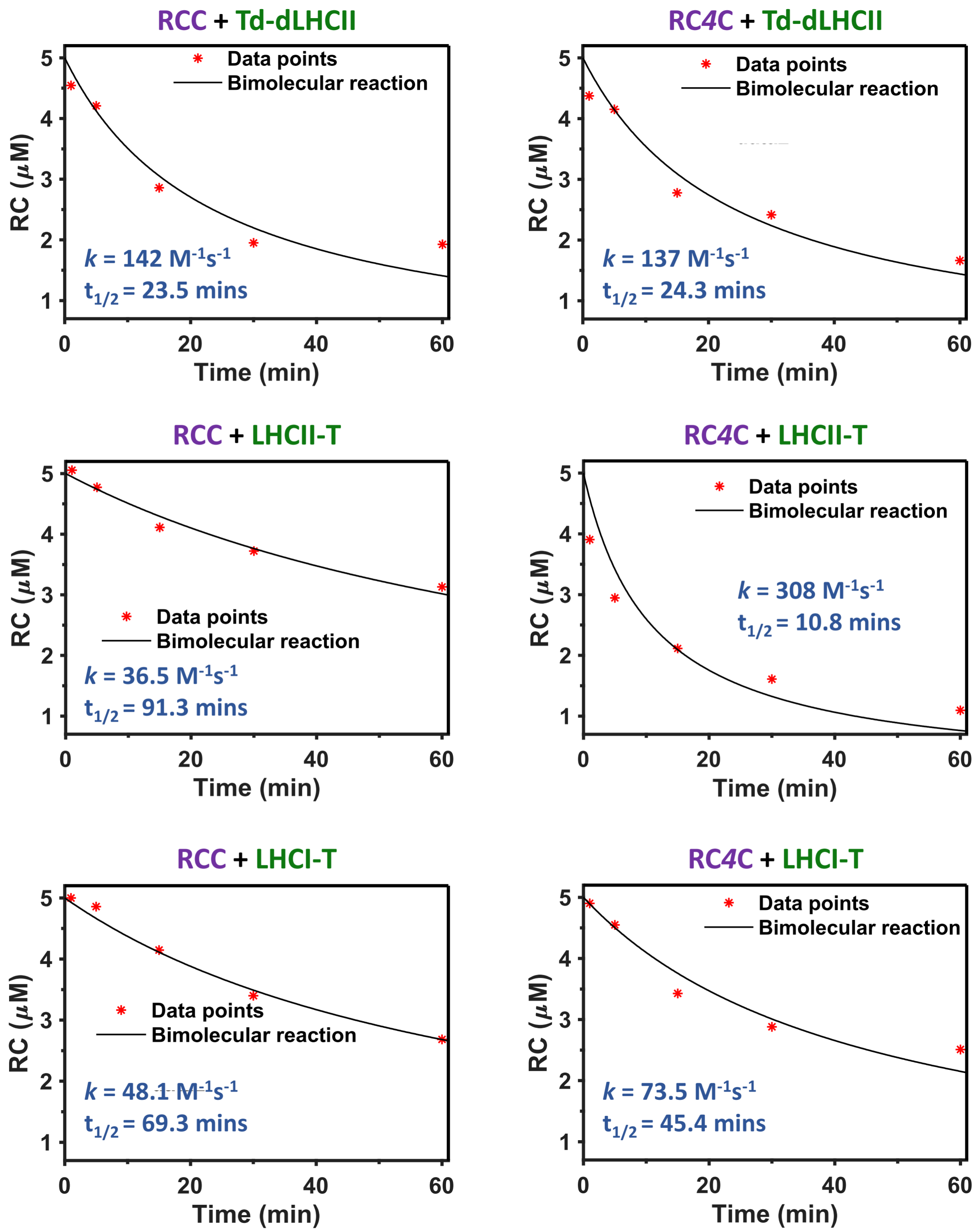

**Supplementary Fig. S19.** Estimation of the rate of chimera formation by SpyTag/SpyCatcher. The rate of chimera formation was quantified from depletion of the SpyCatcherΔ-PufL band in SDS-PAGE gels stained with Coomassie (Fig. S18) and subjected to densitometry (initial protein loading 5 μM). An internal calibration was applied using the RC PuhA or PufM band, depending on which was best distinguished from the LHC bands. Data were fitted to a bimolecular reaction. The reaction constant *k* was then estimated and the half reaction time was deduced based on this reaction constant and an initial concentration of 5 μM, following bimolecular reaction kinetics.

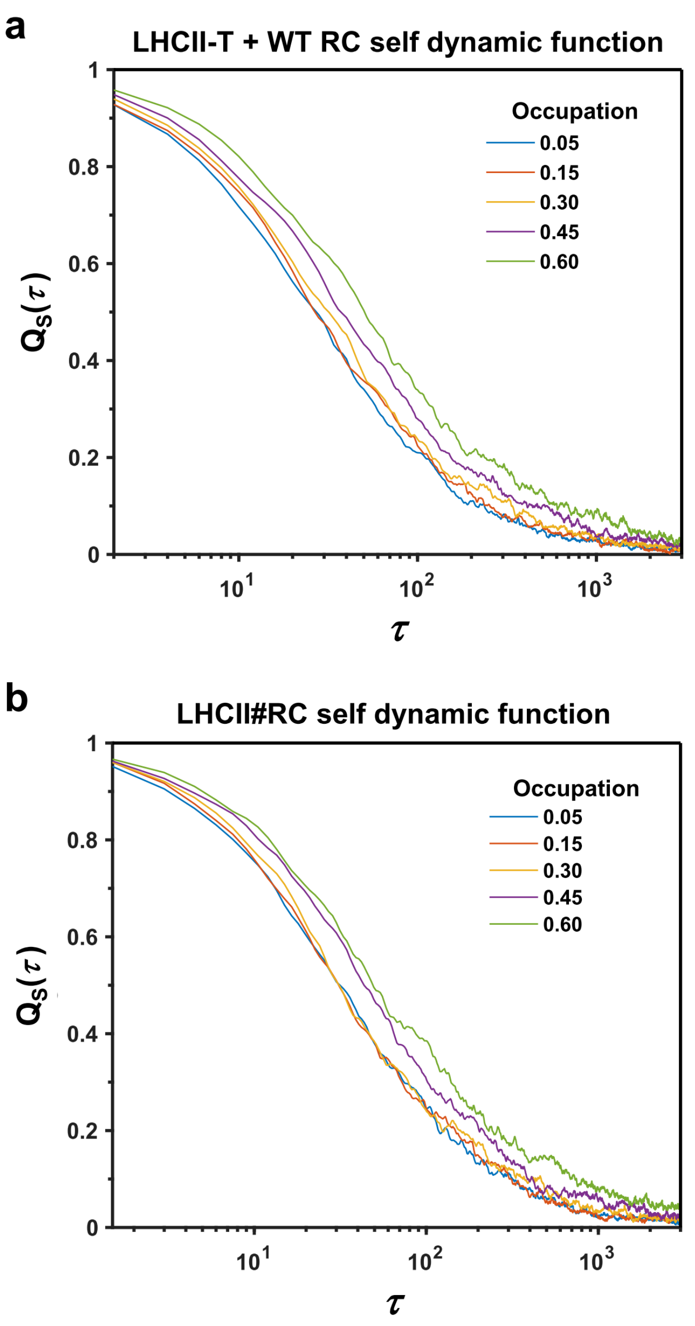

**Supplementary Fig. S20.** Self-dynamic function ($Q_S$) as a function of simulation time ($\tau$). **a** Relaxation of $Q_s$ for a mixture of LHCII-T and WT RC proteins at selected percentage occupations of 5, 15, 30, 45 and 60%. **b** Relaxation of $Q_s$ for LHCII#RC chimeras at the same selected percentage occupations. In both cases $Q_S$ decayed to nearly zero at the end of simulation showing that the system was at equilibrium.

**Supplementary Tables**

**Supplementary Table S1.** Spectral overlap and quantum yield of each system

| System | $J^a$ ($\times10^{-13}$ cm$^{-3}$) | quantum yield[b] (%) |
|---|---|---|
| dLHCII +WT RC | 9.34 | 13.5 |
| Td-dLHCII +WT RC | 9.33 | 13.8 |
| LHCII-T + WT RC | 9.34 | 14.0 |
| LHCI-Td + WT RC | 16.8 | 4.05 |
| Td-LHCI-Td +WT RC | 17.1 | 3.78 |

[a] Spectral overlap of LHC emission with RC molar absorbance, normalised to that of dLHCII and RC (SD < 0.01).

[b] Quantum yield relative to that of dLHCII, obtained by comparing integration of LHC emission (SD in the range 0.01 ~ 0.02).

**Supplementary Table S2.** Nomenclature and descriptions of pigment-protein complexes.

| Name | Description |
|---|---|
| SpyCatcherΔ | SpyCatcher lacking nine C-terminal amino acids |
| SpyTagΔ | SpyTag lacking three C-terminal amino acids |
| RC*C* | SpyCatcherΔ attached to N-terminus of RC PufL |
| RC*4C* | SpyCatcherΔ attached to N-terminus of RC PufL by a 4 amino acid linker |
| dLHCII | Lhcb1 lacking 12 N-terminal amino acids, His tag at C-terminus |
| T-dLHCII | His-tag and full SpyTag added to N-terminus of dLHCII |
| Td-dLHCII | His-tag and truncated SpyTagΔ added to N-terminus of dLHCII |
| LHCII-T | Full SpyTag and His-tag added to C-terminus of Lhcb1 |
| LHCII#RC | Chimera of LHCII-T and RC*C* |
| RC#LHCII | Chimera of RC*4C* and Td-dLHCII |
| Td-L4 | His-tag and SpyTagΔ added to N-terminus of Lhca4 |
| L1 | Myc-tag added to N-terminus of Lhca1 |
| Td-L1 | Myc-tag and SpyTagΔ added to N-terminus of Lhca1 |
| LHCI-Td | LHCI heterodimer from Td-L4 and L1 |
| Td-LHCI-Td | LHCI heterodimer from Td-L4 and Td-L1 |
| LHCI#RC | Chimera of LHCI-Td and one RC*C* |
| RC#LHCI#RC | Chimera of Td-LHCI-Td and two RC*C* |

**Supplementary Table S3.** Summary of fitting of P870 photobleaching and computed photon absorbance.

| Sample | $k_f$ ($10^{-2}$ $s^{-1}$) | $k_r$ ($10^{-2}$ $s^{-1}$) | **RC photon absorbance**[b] ($10^{12}$ $cm^{-2}s^{-1}$) | **LHC photon absorbance**[b] ($10^{12}$ $cm^{-2}s^{-1}$) |
|---|---|---|---|---|
| WT RC | 6.2 ± 0.8 | 79.3 ± 11.7 | 2.0 | - |
| WT RC + dLHCII | 6.9 ± 0.7 | 72.3 ± 8.9 | 2.1 | 28.8 |
| WT RC + LHCII-T | 7.2 ± 0.7 | 67.0 ± 7.7 | 2.0 | 27.7 |
| WT RC + LHCI-Td | 8.0 ±1.6 | 77.9± 19.0 | 1.0 | 23.0 |
| WT RC + Td-LHCI-Td | 8.5 ± 0.9 | 79.9 ± 10.2 | 2.0 | 22.2 |
| RC*C* | 2.6 ± 0.4 | 59.5 ± 15.7 | 2.3 | - |
| RC*C* + dLHCII | 2.5 ± 0.4 | 46.1 ± 11.2 | 1.9 | 26.2 |
| LHCII#RC | 11.3 ± 0.9 | 71.2 ± 7.0 | 1.9 | 27.9 |
| LHCI#RC | 14.3 ± 1.5 | 73.8 ± 9.2 | 1.3 | 28.9 |
| RC#LHCI#RC | 11.1 ± 0.7 | 70.4 ± 5.1 | 3.4 | 38.2 |
| RC*4C* | 2.4 ± 0.4 | 56.9 ± 15.5 | 2.3 | - |
| RC*4C* + dLHCII | 2.7 ± 0.4 | 51.4 ± 10.3 | 2.5 | 26.6 |
| RC#LHCII | 10.1 ± 1.1 | 66.3 ± 8.5 | 1.6 | 22.9 |

[a] Fitted first order kinetic constants ± standard deviation.

[b] Number of photons absorbed by RCs and LHCs per unit area per second calculated from integration across spectra of the product of incident light power and protein absorbance (1 - transmission).

**Supplementary Table S4 .** Estimates of ET efficiency in additional control protein mixtures.

| System | $E_{P870}$ (%) | $E_{FL}$ (%) |
|---|---|---|
| WT RC + dLHCII | 0.8 ± 1.3 | 2.8 ± 1.0 |
| RC*C* + dLHCII | -0.3 ± 1.7 | 1.7 ± 2.5 |
| RC*4C* + dLHCII | 1.2 ± 2.7 | 1.9 ± 1.3 |
